# Role of the NOS-cGMP pathways in time-dependent sensitization of behavioral responses in zebrafish

**DOI:** 10.1101/2024.03.27.586937

**Authors:** Eveline Bezerra de Sousa, João Alphonse Apóstolo Heymbeeck, Leonardo Miranda Feitosa, Amanda Gabriele Oliveira Xavier, Kimberly dos Santos Campos, Lais do Socorro dos Santos Rodrigues, Larissa Mota de Freitas, Rhayra Xavier do Carmo Silva, Saulo Rivera Ikeda, Suellen de Nazaré dos Santos Silva, Sueslene Prado Rocha, Wilker Leite do Nascimento, Edinaldo Rogério da Silva Moraes, Luana Ketlen Reis Leão, Anderson Manoel Herculano, Caio Maximino, Antonio Pereira, Monica Lima-Maximino

**Affiliations:** Laboratório de Bacteriologia e Neuropatologia, Universidade do Estado do Pará – Campus VIII, Marabá/PA, Brazil; Programa de Pós-Graduação em Neurociências e Biologia Celular, Instituto de Estudos em Saúde e Biológicas, Universidade Federal do Pará (UFPA), Belém/PA, Brazil; Laboratório de Neurofarmacologia e Biofísica, Universidade do Estado do Pará – Campus VIII, Marabá/PA, Brazil; Programa de Pós-Graduação em Neurociências e Comportamento, Núcleo de Teoria e Pesquisa do Comportamento, Universidade Federal do Pará (UFPA), Belém/PA, Brazil; Departamento de Morfologia e Ciências Fisiológicas, Universidade do Estado do Pará – Campus VIII, Marabá/PA, Brazil; Departamento de Morfologia e Ciências Fisiológicas, Universidade do Estado do Pará – Campus XXIII, Parauapebas/PA, Brazil; Laboratório de Neurofarmacologia Experimental, Instituto de Ciências Biológicas, Universidade Federal do Pará (UFPA), Belém/PA, Brazil; Laboratório de Neurociências e Comportamento “Frederico Guilherme Graeff”, Instituto de Estudos em Saúde e Biológicas, Universidade Federal do Sul e Sudeste do Pará (Unifesspa), Marabá/PA, Brazil; Laboratório de Processamento de Sinais, Instituto de Tecnologia, Universidade Federal do Pará (UFPA), Belém/PA, Brazil

**Keywords:** Time-dependent sensitization, Stress, Gaseous transmitters

## Abstract

Nitric oxide (NO) is a molecule involved in plasticity across levels and systems. The role of NOergic pathways in time-dependent sensitization, a behavioral model of translational relevance to trauma and stress-related disorders, was assessed in adult zebrafish. In this model, adult zebrafish acutely exposed to a fear-inducing conspecific alarm substance (CAS) and left undisturbed for an incubation period show increased anxiety-like behavior 24 h after exposure. CAS increased forebrain glutamate immediately after stress and 30 min after stress, an effect that was accompanied by increased nitrite levels immediately after stress, 30 min after stress, 90 min after stress, and 24 h after stress. CAS also increased nitrite levels in the head kidney, where cortisol is produced in zebrafish. CAS-elicited nitrite responses in the forebrain 90 min (but not 30 min) after stress were prevented by a NOS-2 blocker. Blocking NOS-1 30 min after stress prevents TDS; blocking NOS-2 90 min after stress also prevents TDS, as does blocking calcium-activated potassium channels in this latter time window. TDS is also prevented by blocking guanylate cyclase activation in both time windows, and cGMP-dependent channel activation in the second time window. These results suggest that different NO-related pathways converge at different time windows of the incubation period to induce TDS.

## 1. Introduction

Time-dependent sensitization (TDS) is a neurobehavioral phenomenon in which behavior is sensitized after an “incubation” period in which the animal (including humans), previously being stressed, is kept stress-free (Stam, 2007b, 2007a; Stam et al., 2000). It represents not simply a sustained response to stressors, but an actual sensitization in which, during the incubation period, associative and non-associative mechanisms contribute to produce increased behavioral and endocrine responses to further stressors (Daskalakis et al., 2013a). TDS is relevant to understand the pathophysiology of delayed and sustained responses to stressors, which in its turn form part of the basis for trauma and stress-related disorders, such as acute stress disorder and post-traumatic stress disorder (PTSD) (Yehuda & Antelman, 1993). In a zebrafish variation, animals are exposed to a relatively serious stressor – the conspecific alarm substance (CAS), which induces fear-like responses (Jesuthasan & Mathuru, 2008; Lima-Maximino et al., 2020; Maximino et al., 2019) - and, after an “incubation” period in which the animal is left undisturbed, assaying anxiety-like behavior in tests such as the light/dark test or the novel tank test (Lima et al., 2016). A variation of this protocol involves a combination of three stressors for 90 min; one week after this complex combined stressor, zebrafish show increased anxiety-like behavior and elevated whole-body cortisol, as well as an upregulation in the expression of brain gene expression of inflammatory molecules (Yang et al., 2020).

Using TDS in zebrafish, it was previously shown that nitric oxide (NO) plays an important role in the consolidation, but not the initiation, of these responses (Lima et al., 2015). Nitric oxide is a gaseous transmitter that has great potential as a mediator of stress sensitization; it has been involved in the consolidation of aversive memories (Akar et al., 2007; A. V. Calixto et al., 2001; Xu et al., 2007a) and unconditioned anxiety responses (Akar et al., 2007; A. V Calixto et al., 2008; Dunn et al., 1998; Herculano et al., 2015; Pokk & Väli, 2002; Spiacci Jr et al., 2008) and was associated with genetic risk factors for PTSD (Bruenig et al., 2017; Lawford et al., 2013).

NO is the product of the transformation of arginine into citrulline by nitric oxide synthase (NOS), an isozyme with three different isoforms (Garthwaite, 2008). Two isoforms, NOS-1 and NOS-3, are *constitutive* and calcium-dependent, while a third isoform, NOS-2, is inducible and calcium-independent (Mittal & Kakkar, 2020). Since NOS-2 is not constitutively expressed in normal, healthy organisms, its activity is usually delayed, and involves novel protein transcription that is initiated by stressful and/or inflammatory stimuli (Thomas et al., 2015); moreover, since it does not depend on calcium, its activity is much more prolonged, depending on NO clearance by diffusion, enzyme breakdown, or metabolism of downstream mediators by phosphodiesterases to stop producing NO (Thomas et al., 2015). One source of NOS-2 upregulation and activation is calcium-activated potassium (KCNN) channels in microglia (Kaushal et al., 2007). From the constitutive isoforms, NOS-1 is perhaps best know for its close relation with the glutamatergic NMDA receptor (NMDA-R) (Luo & Zhu, 2011): after depolarization and opening of a magnesium gate, activation of this receptor leads to a calcium influx and, due to the physical connection with NOS-1 (Kornau et al., 1995), to the activation of this isoform and consequent production of NO. The “canonical” signaling pathway for NOS isozymes is dependent on the activation of guanylate cyclase and consequent production of cyclic guanosine monophosphate (cGMP), although non-canonical mechanisms have also been described (Calabrese et al., 2007; Garthwaite, 2008). This mechanism has been implicated in neuroplasticity (Edwards & Rickard, 2007), while NOS-2 is usually associated with neurotoxicity and neuroinflammation (Calabrese et al., 2007). Both processes are thought to mediate stress sensitization (Adamec et al., 1998, 2001, 2006; Baker et al., 2012; Harvey, 2006).

One putative mechanism for stress sensitization mediated by nitric oxide is metaplasticity, the persistent neuronal responses elicited synaptic activity that are not manifested as changes in synaptic strength, leading to concomitant changes in the neuron’s ability to generate synaptic plasticity (Abraham & Tate, 1997). After activation of NMDA-Rs, the production of NO can provide a negative feedback mechanism, elevating the threshold for long-term potentiation (Abraham, 2008). Metaplasticity can support adaptive changes during distinct stages of the formation of aversive memories, through either associative and non-associative mechanisms, in brain region-specific manners (Çalışkan & Stork, 2018). The term “behavioral metaplasticity” has been introduced to reflect the fact that environmental stimuli can act as “triggers” or “primers” to promote long-term changes in both synaptic metaplasticity and at the behavioral level (Abraham & Richter-Levin, 2018). Particularly, environmental stressors can influence the capacity for long-term synaptic plasticity in limbic structures, either through circulating glucocorticoids or other mechanisms (Schmidt et al., 2013). Indeed, in a rodent model of predator exposure, stress induces long-lasting potentiation in an input to the basolateral amygdala, leading to increased anxiety-like behavior 9 days after exposure (Adamec et al., 2005a). This effect is prevented by blocking NMDA-Rs (Adamec et al., 2005b).

In a rodent model of PTSD that is conceptually related to TDS, repeated stress caused increases in hippocampal nitrite levels – indicative of increased NO production – on day 7 post-stress, an effect that was blocked by the NOS-1 inhibitor 7-nitroindazole (7-NI) (Harvey et al., 2005). 21 days after the application of TDS, hippocampal NOS activity was increased, an effect that was blocked by the NOS-2 inhibitor aminoguanidine (AG), but not 7-NI (Harvey et al., 2004). These results suggest that both NOS-1 and NOS-2 participate in the production of NO in time-dependent sensitization, but at different time intervals. In the single-prolonged stress model, conditional knockout of NOS-1 in the dorsal raphe nucleus of mice rescues the PTSD-like phenotype (increased anxiety-like behavior, enhanced contextual fear memory, and fear generalization) (Sadeghi et al., 2022). A participation of NOS isozymes in the behavioral effects of TDS was demonstrated in zebrafish, in which treatment with the non-selective inhibitor L-NAME 30 and 90 min. after CAS exposure blocks the development of increased anxiety 24 h after exposure (Lima et al., 2015). Currently, it is not known which isoform is responsible for the behavioral effects of TDS in zebrafish.

In the present work, we tested the hypothesis that CAS would lead to the activation of NOS-1 in the limbic telencephalon due to an glutamate efflux immediately after stress, and that, at a posterior time window of the incubation period, synthesis and activation of NOS-2 would lead to a sustained production of NO. Both processes are hypothesized to contribute to the increased anxiety-like behavior that is observed 24 h after CAS exposure. Moreover, we hypothesized that cGMP would also mediated the effects of TDS at both time windows. Finally, we hypothesized that blocking KCNN channels would lead to the inhibition of TDS

## 2. Methods

### 2.1. Reagents

All drugs were bough from Sigma-Aldrich, as follows: aminoguanidine hydrochloride (AG): CAS #1937-19-5, Sigma-Aldrich 396494; 7-Nitro-1*H*-indazole (7-nitroindazole, 7-NI): CAS# 2942-42-9, Sigma-Aldrich N7778; triarylmethane-34 (1-[(2-Chlorophenyl)diphenylmethyl]-1H-pyrazole, TRAM-34): CAS #289905-88-0, Sigma-Aldrich T6700; 1H-[1,2,4]Oxadiazolo[4,3-a]quinoxalin-1-one (ODQ): CAS #41443-28-1, Sigma-Aldrich O3636; 3-[3-({[(7S)-3,4-dimethoxybicyclo[4.2.0]octa-1,3,5-trien-7-yl]methyl}(methyl)amino)propyl]-7,8-dimethoxy-2,3,4,5-tetrahydro-1H-3-benzazepin-2-one (ivabradine hydrochloride): CAS #148849-67-6, Sigma-Aldrich SML0281).

Reagents for the glutamate assay were obtained from Sigma-Aldrich, as follows: methanol: CAS #67-56-1, Sigma-Aldrich 439193; 2-propanol: CAS #67-63-0, Sigma-Aldrich 34863; sodium acetate: CAS #127-09-3, Sigma-Aldrich 241245; boric acid: CAS #10043-35-3, Sigma-Aldrich B0394; zinc chloride: CAS #7646-85-7, Sigma-Aldrich 208086; *o*-phtalaldehyde (OPA): CAS #643-79-8, Sigma-Aldrich P0532; glutamate standard: CAS #56-86-0, Sigma-Aldrich G1251; homoserine standard: CAS #672-15-1, Sigma-Aldrich H6515. All reagents were of the highest grade available, and HPLC-grade whenever possible.

Reagents for the nitrite assay (sodium nitrite standard [CAS # 7632-00-0]: Sigma-Aldrich 237213, ACS reagent, ≥97.0%; sulfanilamide [CAS #63-74-1]: Sigma-Aldrich S9251, ≥98.0%; *N*-(1-naphthyl)ethylenediamine [CAS #1465-25-4]: Sigma-Aldrich 222488 ACS reagent, ≥98.0%) were donated by Dr. Valney Mara Gomes Conde (https://www.researchgate.net/profile/Valney-Conde-2).

### 2.2. Animals and housing

1064 adult zebrafish from the longfin phenotype were used in the present experiments. Animals were group-housed in mixed-sex 40 L glass tanks (maximum density of 25 fish per tank) during acclimation, with an approximate ratio of 1:1 males to females (estimated by body morphology). Animals were acquired from commercial vendors (Fernando Peixes, Belém/PA, and PiciculturaPower Fish, Itaguaí/RJ), and delivered to the laboratory with an approximate age of 3 months (standard length = 12.9 ± 1.6 mm), being quarantined for at least two weeks before experiments begun. Throughout experiments, animals were estimated to be 4 months old (standard length = 21.7 ± 3.1 mm). Outbred populations were used to their higher genetic variability, which is expected to decrease the impacts of random genetic drift that could lead to the development of uniquely heritable neurobehavioral traits (Parra et al., 2009; Speedie & Gerlai, 2008).These populations are expected to better represent the natural populations in the wild, and have previously been shown to respond to conspecific alarm substance (Lima et al., 2015, 2016; Speedie & Gerlai, 2008). Breeders were licensed for aquaculture under Ibama’s (Instituto Brasileiro do Meio Ambiente e dos Recursos Naturais Renováveis) Resolution 95/1993.

Acclimation tanks were filled with non-chlorinated water kept at room temperature (28 °C), and a pH of 7.0-8.0. Lighting was provided in a cycle of 14 hours-10 hours of darkness by fluorescent lamps in the ceiling, according to standards of care for zebrafish (Lawrence, 2007). Other water quality parameters were as follows: hardness 100-150 mg/L CaCO3; dissolved oxygen 7.5-8.0 mg/L; ammonia and nitrite < 0.001 ppm. Potential suffering of animals was minimized by controlling for the aforementioned environmental variables and scoring humane endpoints (clinical signs, behavioral changes, bacteriological status), following Brazilian legislation (Conselho Nacional de Controle de Experimentação Animal - CONCEA, 2017). Animals were used for only one experiment and in a single behavioral test, to reduce interference from apparatus exposure. Experiments were approved by UEPA’s IACUC under protocol 06/18.

### 2.3. CAS preparation and administration

CAS was extracted from the skin of conspecific donors using the protocol at https://dx.doi.org/10.17504/protocols.io.tr3em8n. Animals were exposed in groups of 5 individuals with CAS extracted from one animal, in a concentration of 7 mL of CAS in 5 L of water. Exposure was made for 6 min., after which animals were removed from the exposure tank and left to rest for 24 h, in the same group of 5 individuals. Control animals were exposed to the same volume of distilled water. Experimenters were blinded to treatment by using coded vials.

### 2.4. Forebrain dissection and glutamate quantification

To quantify glutamate levels in the ECF, animals were sacrificed at different time points after CAS exposure (immediately after exposure [0 min.], 15 min., 30 min., 90 min. and 24 h after exposure; Figure 1A), and their brains were quickly removed and incubated in extraction solution, following the protocol describe in dx.doi.org/10.17504/protocols.io.14egn3p1zl5d/v1.

**Figure 1.**
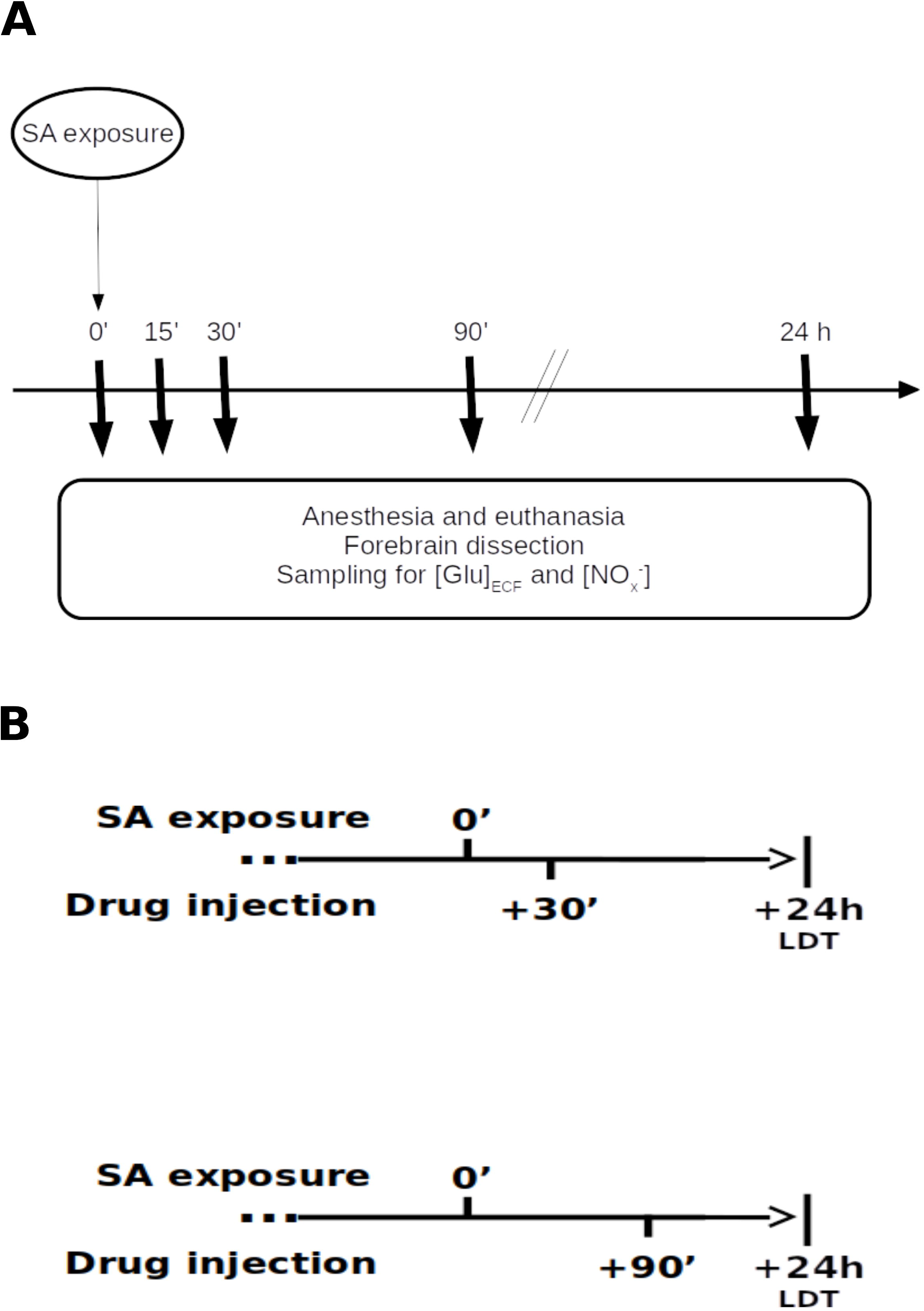
Experimental designs for (A) neurochemical experiments and (B) behavioral experiments. In (A), animals are exposed to CAS or water at time = 0’, and sacrificed at different time intervals (0’, 15’, 30’, 90’, and 24 h after exposure). In (B), animals are exposed to CAS or water at time = 0’, injected with vehicle or drug at 30’ or 90’, and left undisturbed for 24 h before the light/dark preference test.

Glutamate levels in samples were assayed using fluorescent high-performance liquid chromatography (HPLC) through a method previously used to quantify aminoacids in ECF of retinal explants (Moraes et al., 2012). A Shimadzu HPLC (LC20-AT model) was used, with RF-10Axl fluorescence detector, DGA20A5 degasser, CBM20A communication module, Rheodyne injection system with a 20 μL injection loop, and Shim-Pack VP-ODS chromatography column (250 x 4.6 mm dimension, with 5 μm particles). Samples were injected in the column for separation using a Hamilton 50 μL microsyringe. Before injection, a pre-column derivatization agent, *o-* phtalaldehyde (OPA), was added as a flurophore, in a solution of OPA (13 mg) and *N*-acetilcysteine (16.3 mg) in 15% methanol, as well as a solution of borate buffer (pH 9.5). The solutions were added to the sample in a proportion of 1 volume of sample:6 volumes of OPA solution:4 volumes of buffer solution. The derivatization reaction, completed after 5 min. of incubation, forms a isoindol fluorescent indole of aminoacids in the sample.

After derivatization, samples were injected in the HPLC system, using reverse chromatography with elution gradient with two mobile phases. Phase A was composed of sodium acetate buffer 50 μM, 5% methanol, and 12 mL propanol for every liter of phase. Phase B was composed of 70% methanol. A gradient was produced with 100% phase A from 0 to 10 min. of the assay, 70% phase A and 30% phase B from 10 min to 20 min of the assay, and 100% phase A from 25 min onwards. The elution flow speed was 1.2 mL/min.

After separation in the reverse HPLC system, internal standard and glutamate contained in samples were detected and quantified by a fluorescence detector, with excitation wavelength of 340 nm and emission wavelength of 460 nm. The concentration of glutamate was calculated via the ratio of the height of the peak identified in the chromatogram and the homoserine peak. All analyses were made in triplicate.

### 2.5. Quantification of nitrite levels in brains and head kidneys

Brains were dissected as above, and head kidneys were removed and dissected, following the protocol at Gerlach et al. (2011). Nitrite levels were assessed using an optimized Griess (1864) assay, as described in the protocol at https://dx.doi.org/10.17504/protocols.io.sabeaan.

### 2.6. Drug injection

For drug injection, the protocol described by Kinkel et al. (2010) was followed. Briefly, 30 minutes or 90 minutes after exposure, animals were anesthetized in ice-cold water (temperature between 12 °C and 14 °C), transferred to a water-soaked sponge surgical bed, and injected with the drug or vehicle, using a microsyringe (Hamilton® 701 N syringe, needle size 26 gauge at cone tip), with total volumes of injection of 5 μL. Drug doses (Table 1) were based on data from rodent literature. The time intervals for injection after CAS exposure were based on previous data which showed that treatment with L-NAME in these intervals prevented the development of TDS in zebrafish (Lima et al., 2015).

**Table 1.** Drug targets and doses used in the experiments.

| Drug | Target | Dose | Reference |
| --- | --- | --- | --- |
| 7-NI | NOS-1 | 30 mg/kg | MacKenzie et al., 1994 |
| Aminoguanidine | NOS-2 | 50 mg/kg | da Silva Chaves et al., 2020 |
| TRAM-34 | KCNN channels | 10 mg/kg | Chen et al., 2011 |
| ODQ | Soluble guanylate cyclase | 10 mg/kg | Mutlu et al., 2015 |
| Ivabradine | Nucleotide-gated channels | 15 mg/kg | Łuszczki et al., 2013 |

### 2.5. Light-dark test

24 h after CAS, animals were tested in the light-dark test (LDT), an assay for anxiety-like behavior (Maximino et al., 2010). The test was run according to the protocol at https://dx.doi.org/10.17504/protocols.io.srfed3n. Video files were analyzed using X-Plo-Rat, and the following variables were coded and extracted:

- Time spent in the white compartment (s), the main endpoint
- Entries in the white compartment (N)
- Average duration of entries on white (s), calculated by dividing the time spent in the compartment by the number of entries
- Risk assessment (N), defined as the number of entries with duration lower than 1 s, or partial entries, in which the individual crosses only half of its body in the tank midline
- Erratic swimming (N), defined as the number of entries in the white compartment with fast, highly angular swimming with unpredictable trajectory
- Thigmotaxis (%), the percentage of time in the white compartment spent in less tan 2 cm from one tank wall.

### 2.8. Statistical analysis

For experiments on the time-course of neurochemical changes in the forebrain, data were analyzed using a mixed analysis of variance (within-subjects factor: time point; between-subjects factor: treatment). For behavioral experiments, as well as for the experiment on the role of NOS-2 on nitrite elevations, data were analyzed using two-way (treatment X drug) analysis of variances.

## 3. Results

### 3.1. Neurochemical analyses

#### 3.1.1. Forebrain glutamate efflux is increased immediately after CAS exposure

Extracellular levels of glutamate in the forebrain were assessed by immersing dissected forebrains in an extraction fluid (Maximino et al., 2013), followed by HPLC measurement (Moraes et al., 2012). Two glutamate peaks were observed in animals exposed to CAS (exposure: F_[1, 40]_ = 8.19, p = 0.0007; time after exposure: F_[4, 40]_ = 3.11, p = 0.026; interaction: F_[4, 40]_ = 6.94, p = 0.008), with increases observed in animals sacrificed immediately after CAS and 30 min after CAS (Figure 2A).

**Figure 2.**
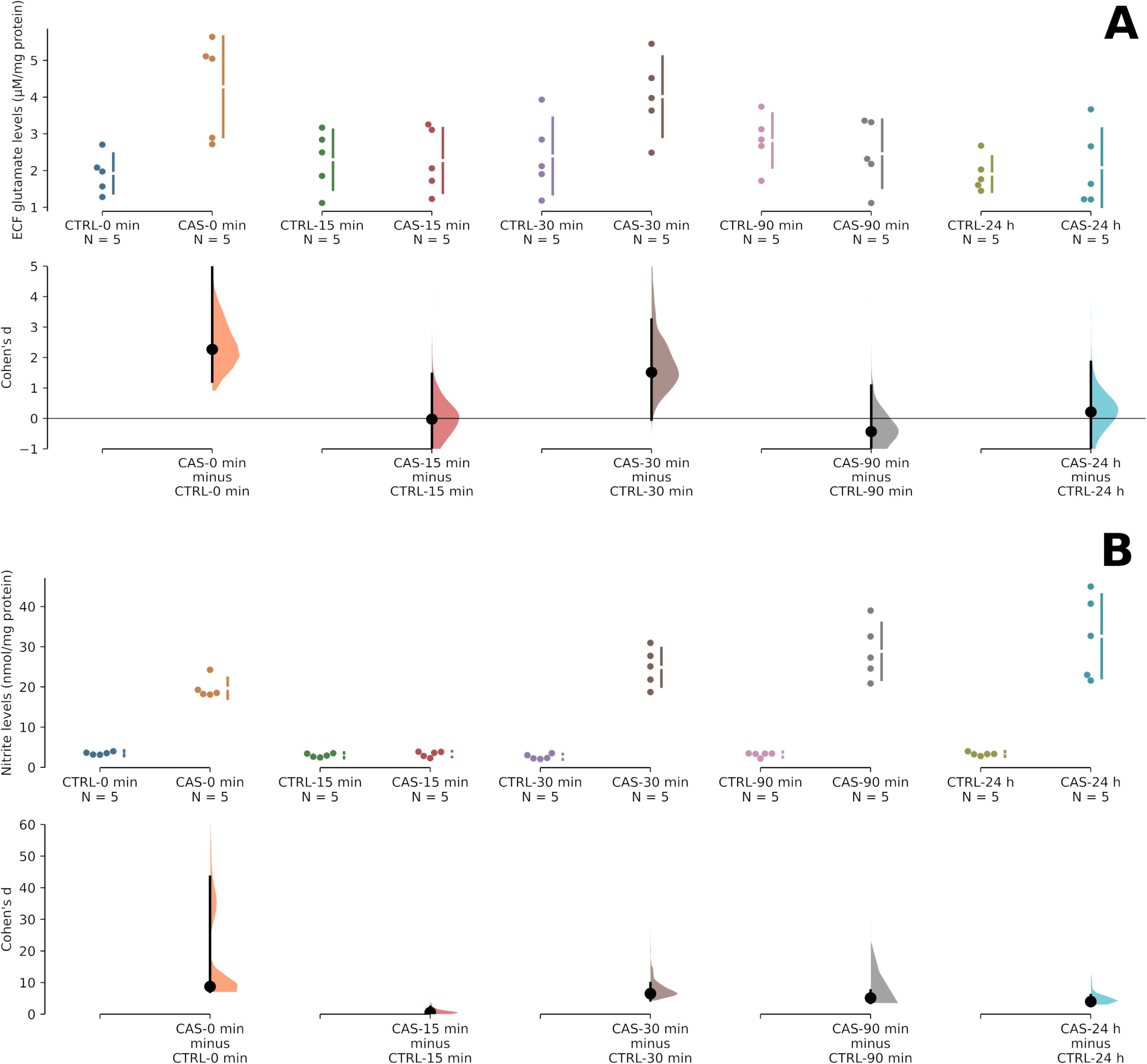
CAS increases ECF glutamate in the zebrafish forebrain immediately and 30 minutes after exposure, and increases tissue nitrite levels immediately after stress, and 30 min, 90 min, and 24 h after stress. (A) Concentrations of glutamate in the extracellular fluid of zebrafish forebrains at different time intervals after exposure to water (CTRL) or CAS. (B) Concentrations of nitrite in zebrafish forebrain tissue homogenates at different time intervals after exposure to water or CAS. The raw data is plotted on the upper axes; individual points represent single datapoints (one animal in (A), a pool of five forebrains in (B)). Each effect size (Cohen’s *d*) is plotted on the lower axes as a bootstrap sampling distribution. 5000 bootstrap samples were taken; the confidence interval is bias-corrected and accelerated. Mean differences are depicted as dots; 95% confidence intervals are indicated by the ends of the vertical error bars.

#### 3.1.2. Nitrite levels increase up to 24 h after CAS exposure

The time-course of tissue nitrite (NO_x_^-^) levels in forebrains was assessed in homogenized tissue by the Griess (1864) method. Peaks were observed immediately after exposure, 30 min after exposure, 90 min after exposure, and 24 h after exposure (exposure: F_[1, 40]_ = 230.89, p < 0.001; time: F_[4, 40]_ = 17.4, p < 0.001; interaction: F_[4, 35]_ = 17.03, p < 0.001; Figure 2B).

#### 3.1.3. Participation of NOS-2 in elevated nitrite after CAS exposure

Given the participation of NO on time-dependent sensitization (Lima et al., 2015) and the persistently elevated forebrain nitrite levels observed in the present experiments (Figure 2B), and given the observation that NOS-2 is an inducible enzyme that catalyzes sustained NO production (Thomas et al., 2015), we hypothesized that increased NO_X_^-^ levels 24 h after CAS would be due to NOS-2 activity initiated during the incubation period. NOS-2 was blocked with aminoguanidine (AG, 50 mg/kg) and animals sacrificed either 30 min after exposure or 90 min after exposure. In animals sacrificed 30 min after exposure, a main effect of treatment (F_[1, 6]_ = 7.63, p = 0.033) was found, but no main effect drug (F_[1, 6]_ = 0.11, p = 0.75) was found. A treatment X drug interaction was also absent (F_[1, 6]_ = 0.01, p = 0.913), indicating that AG was unable to block the effect of CAS on the 30 min NO_X_^-^ peak in forebrain samples (Figure 3A). In animals sacrificed 90 min after exposure, main effects of treatment (F_[1, 6]_ = 22.42, p = 0.002) and drug (F_[1, 6]_ = 12.83, p = 0.009) were found, and a treatment X drug interaction was present as well (F_[1, 6]_ = 10.14, p = 0.015). Post-hoc tests revealed differences between animals treated with vehicle and exposed to ddH2O and animals treated with vehicle and exposed to CAS (p = 0.005); and animals treated with vehicle and exposed to CAS and animals treated with AG and exposed to CAS (p = 0.006), indicating that AG was able to block the effects of CAS on the 90 min NO_x_^-^ peak (Figure 3B).

**Figure 3.**
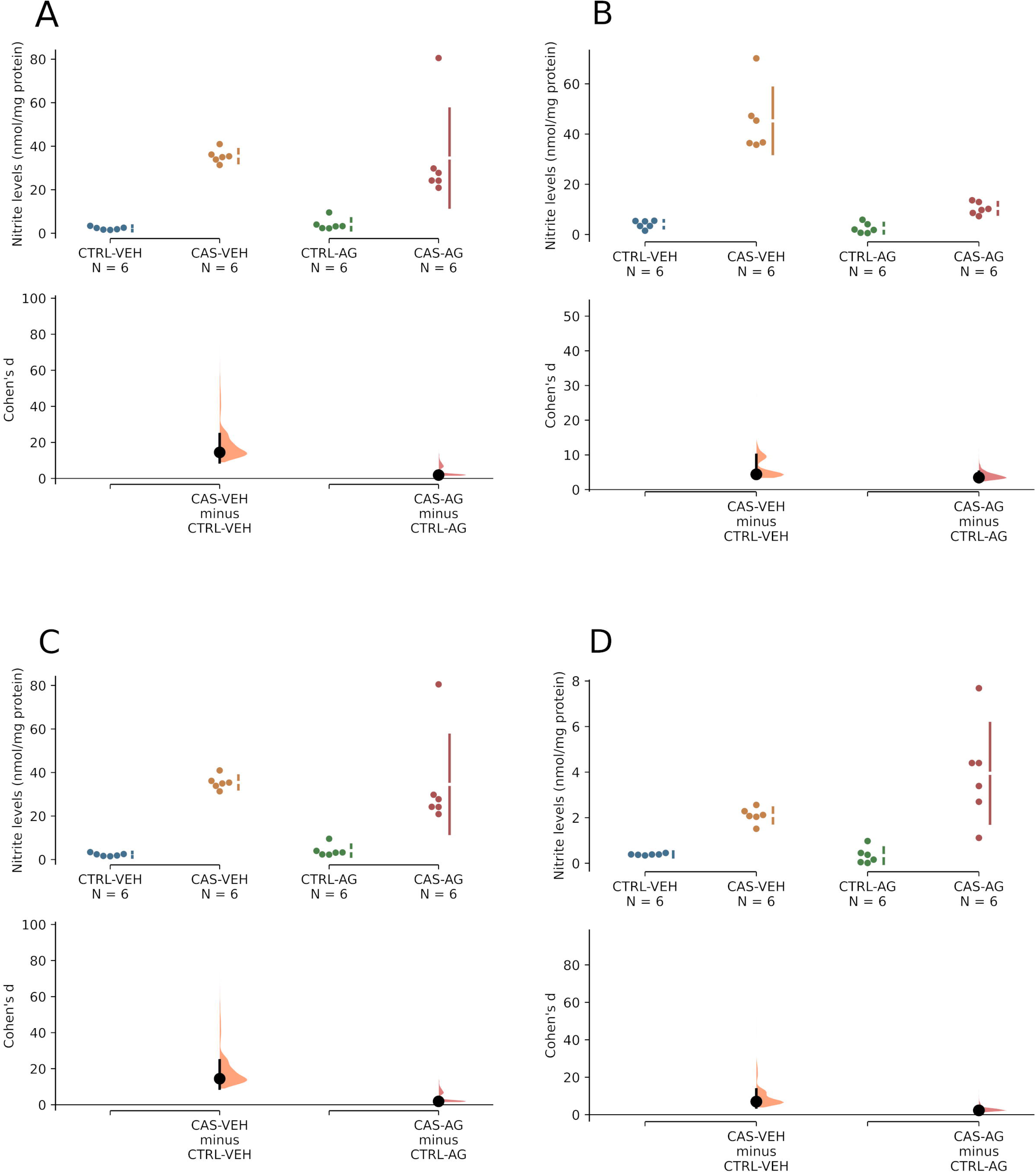
CAS-elicited increases in forebrain (but not head kidney) nitrite levels are mediated by NOS-2 activated 90 minutes (but not 30 minutes) after stress. (A) Forebrain nitrite level 30 minutes after exposure to water (CTRL) or CAS, in animals treated with vehicle (VEH) or 50 mg/kg aminoguanidine (AG). (B) Forebrain nitrite level 90 minutes after exposure to water (CTRL) or CAS, in animals treated with vehicle (VEH) or 50 mg/kg aminoguanidine (AG). (C) Head kidney nitrite level 30 minutes after exposure to water (CTRL) or CAS, in animals treated with vehicle (VEH) or 50 mg/kg aminoguanidine (AG). (D) Forebrain nitrite level 30 minutes after exposure to water (CTRL) or CAS, in animals treated with vehicle (VEH) or 50 mg/kg aminoguanidine (AG). The raw data is plotted on the upper axes; individual points represent individual animals. Each effect size (Cohen’s *d*) is plotted on the lower axes as a bootstrap sampling distribution. 5000 bootstrap samples were taken; the confidence interval is bias-corrected and accelerated. Mean differences are depicted as dots; 95% confidence intervals are indicated by the ends of the vertical error bars.

We also analyzed NO_X_^-^ levels in the head kidney at both time points, with or without AG treatment. The head kidney of teleosts contain interrenal cells, composed of steroidogenic and chromaffin cells, the functional equivalents of the mammalian adrenal cortex and medulla, respectively (Bacila et al., 2021). In animals sacrificed 30 min after exposure, a main effect of treatment (F_[1, 6]_ = 8.58, p = 0.026) was found, but no main effect drug (F_[1, 6]_ = 0.01, p = 0.91) was found. A treatment X drug interaction was also absent (F_[1, 6]_ = 0.05, p = 0.831), indicating that AG was unable to block the effect of CAS on the 30 min NO_X_^-^ peak in head kidney samples (Figure 3C). In animals sacrificed 90 min after exposure, a main effect of treatment (F_[1, 6]_ = 8.54, p = 0.019) was found, but no main effect drug (F_[1, 6]_ = 1.45, p = 0.26) was found. A treatment X drug interaction was also absent (F_[1, 6]_ = 1.3, p = 0.287), indicating that AG was unable to block the effect of CAS on the 90 min NO_X_^-^ peak in head kidney samples (Figure 3D). Overall, these results indicate that CAS elevates nitrite levels 30 and 90 min after exposure in both telencephalon and head kidney, and NOS-2 participates only in the 90 min peak in telencephalic tissue samples.

### 3.2. Behavioral analyses

#### 3.2.1. NOS-1 participates in the first time window of behavioral sensitization: 30 min. after CAS exposure

The elevation of ECF glutamate in the telencephalon immediately after CAS exposure (Fig. 2A) and the fact that AG was unable to block the elevation of NO_X_^-^ 30 min after stress (Fig. 3A), together with the observation that the nonspecific NOS blocker L-NAME blocked the development of TDS when injected 30 and 90 min after CAS (Lima et al., 2015), suggested a participation of NOS-1 in the first time window of behavioral sensitization (30 min after exposure). Given its physical anchoring to NMDA-Rs via the PSD-95 scaffold protein (Cui et al., 2007; Kornau et al., 1995), activation of these receptors by glutamate is thought to be the main trigger of NOS-1 activity. Animals were treated with 7-nitroindazole (7-NI), a specific NOS-1 blocker, 30 min after CAS, and their behavior in the LDT was observed 24 h after stress. A main effect of treatment (F _[1, 36]_ = 10.387, p = 0.0027) was found for time on white (Fig. 4A), but a main effect of drug was absent (F_[1, 36]_ = 1.575, p = 0.2176); a drug X treatment effect was found for this variable (F_[1, 36]_ = 6.206, p = 0.0175). Post-hoc analysis (Tukey’s HSD) found that CAS decreased time on white (p = 0.0016, VEH + CTRL vs. VEH + CAS), an effect that was partially blocked by 7-NI (p = 0.05, VEH + CAS vs. 7-NI + CAS). No main or interaction effects were found for entries on white (treatment: F_[1, 36]_ = 0.257, p = 0.6153; drug: F_[1, 36]_ = 0.011, p = 0.9179; interaction: F_[1, 36]_ = 3.611, p = 0.0654; Figure 4B). No main or interaction effects were found for average duration of entries on white (treatment: F_[1, 36]_ = 1.162, p = 0.288; drug: F_[1, 36]_ = 0.027, p = 0.871; interaction: F_[1, 36]_ = 2.629, p = 0.114; Figure 4C). No main effects were found for risk assessment (treatment: F _[1, 36]_ = 0.717, p = 0.4026; drug: F_[1, 36]_ = 0.549, p = 0.4636), but an interaction effect was found (F_[1,36]_ = 6.634, p = 0.0143; Figure 4D). Post-hoc analysis found that CAS increased risk assessment (p = 0.0406, VEH + CTRL vs. VEH + CAS), an effect that was blocked by 7-NI (p = 0.106, VEH + CAS vs. 7-NI + CAS). No main or interaction effects were found for erratic swimming (treatment: F_[1, 36]_ = 0.14 p = 0.71; drug: F_[1, 36]_ = 0.204, p = 0.654; interaction: F_[1, 36]_ = 2.383, p = 0.131; Figure 4E). Finally, no main or interaction effects were found for thigmotaxis (treatment: F_[1, 36]_ = 2.553, p = 0.127; drug: F_[1, 36]_ = 0.3.185, p = 0.083; interaction: F_[1, 36]_ = 0.268, p = 0.608; Figure 4F). These results suggest that NOS-1 participates in the incubation of TDS in the early periods post-stress.

**Figure 4.**
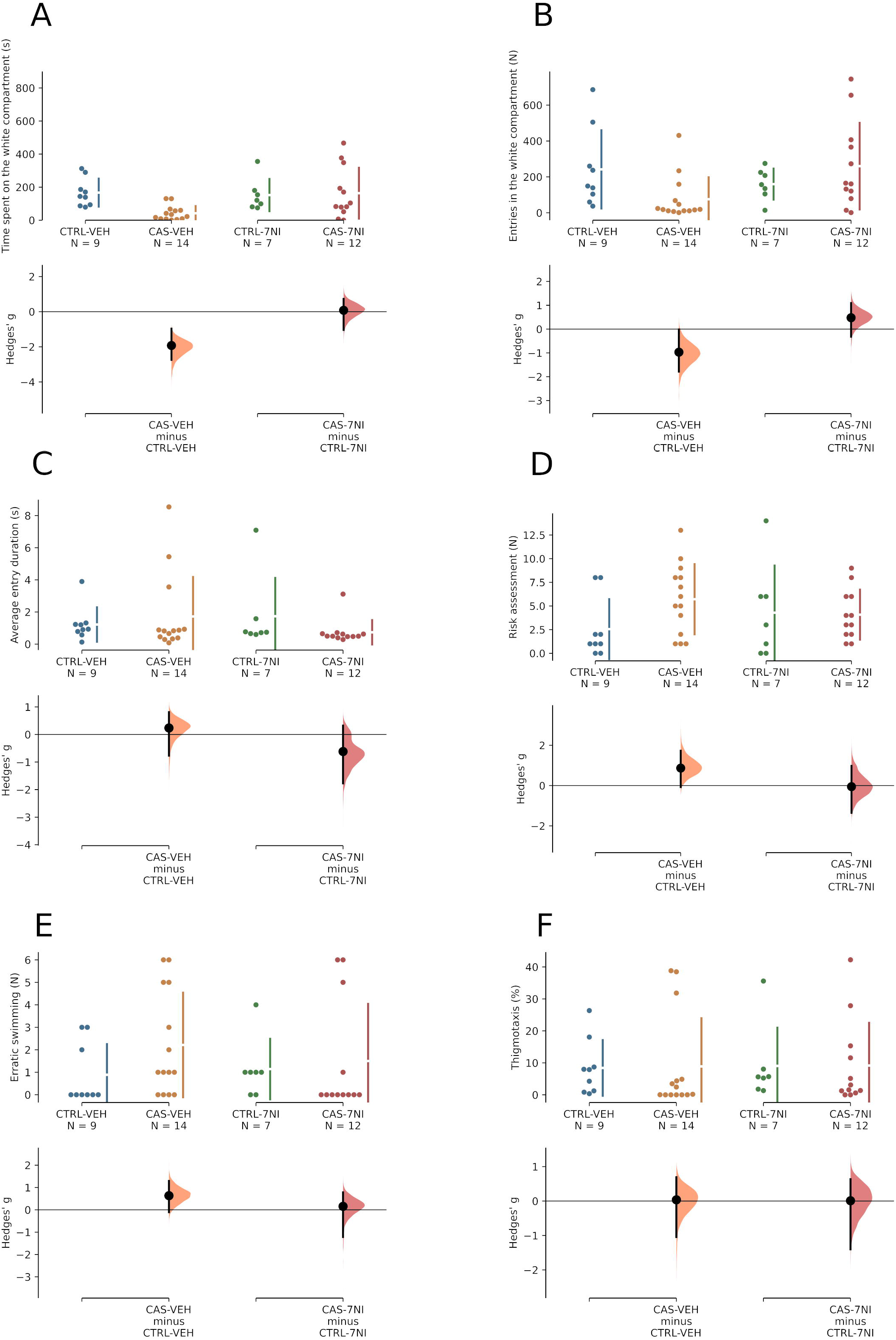
CAS-elicited increases in anxiety-like behavior in the light/dark test 24 h after exposure are mediated by NOS-1 activated 30 minutes after stress. (A) Scototaxis, indexed as time spent in the white compartment. (B) Total locomotion, indexed as total number of entries in the white compartment. (C) Average entry duration. (D) Total number of risk assessment events. (E) Total number of erratic swimming events. (F) Thigmotaxis, indexed by proportion of time in the white compartment spent near the walls. The raw data is plotted on the upper axes; individual points represent individual animals. Each effect size (Cohen’s *d*) is plotted on the lower axes as a bootstrap sampling distribution. 5000 bootstrap samples were taken; the confidence interval is bias-corrected and accelerated. Mean differences are depicted as dots; 95% confidence intervals are indicated by the ends of the vertical error bars.

#### 3.2.2. NOS-2 participates in the second time window of behavioral sensitization: 90 min. after CAS exposure

Sensitization processes can depend on the sustained production of mediators; in the case of NO, the main mechanism through which its production is sustained is through the inducible isoform, NOS-2 (Cinelli et al., 2020). In order to understand the role of this isoform, animals were treated with aminoguanidine (AG), a specific NOS-2 blocker, 30 min. or 90 min. after CAS, and their behavior in the LDT was observed 24 h after stress. When animals were treated with AG 30 min. after CAS, a significant main effect of treatment (F_[1, 34]_ = 12.27, p = 0.001), but not drug (F_[1, 34]_ = 1.42, p = 0.241) was found for time on white; no interaction effect was found (F_[1, 34]_ = 1.66, p = 0.206). Animals exposed to CAS showed decreased time on white 24 h after exposure, an effect that was not blocked by AG (Figure 5A). No significant main or interaction effects were found for entries on the white compartment (treatment: F_[1, 34]_ = 0.77, p = 0.386; drug: F[1, 34] = 0.77, p = 0.385; interaction: F_[1, 34]_ = 1.02, p = 0.319; Figure 5B) or average entry duration (treatment: F_[1, 34]_ = 0.33, p = 0.569; drug: F_[1, 34]_ = 1.42, p = 0.242; interaction: F_[1, 34]_ = 1.47, p = 0.233; Figure 5C). A significant main effect of treatment (F_[1, 34]_ = 4.72, p = 0.0037), but not drug (F_[1, 34]_ = 0.23, p = 0.633) was found for risk assessment; no interaction effect was found (F_[1, 34]_ = 0.3, p = 0.586). Animals exposed to CAS showed increased risk assessment 24 h after exposure, an effect that was not blocked by AG (Figure 5D). No main or interaction effects were found for erratic swimming (treatment: F_[1, 34]_ = 2.12, p = 0.155; drug: F_[1, 34]_ = 3.95, p = 0.055; interaction: F_[1, 34]_ = 0.12, p = 0.731; Figure 5E). A main effect of treatment (F_[1, 34]_= 53.37, p < 0.001), but not of drug (F_[1, 34]_ = 0.06, p = 0.802), was found for thigmotaxis; an interaction effect was not found (F[1, 34] = 0.03, p = 0.856). Animals exposed to CAS showed increased thigmotaxis 24 h after exposure, an effect that was not blocked by AG (Figure 5F). Overall, NOS-2 does not appear to be involved in TDS at the 30 min. time window.

**Figure 5.**
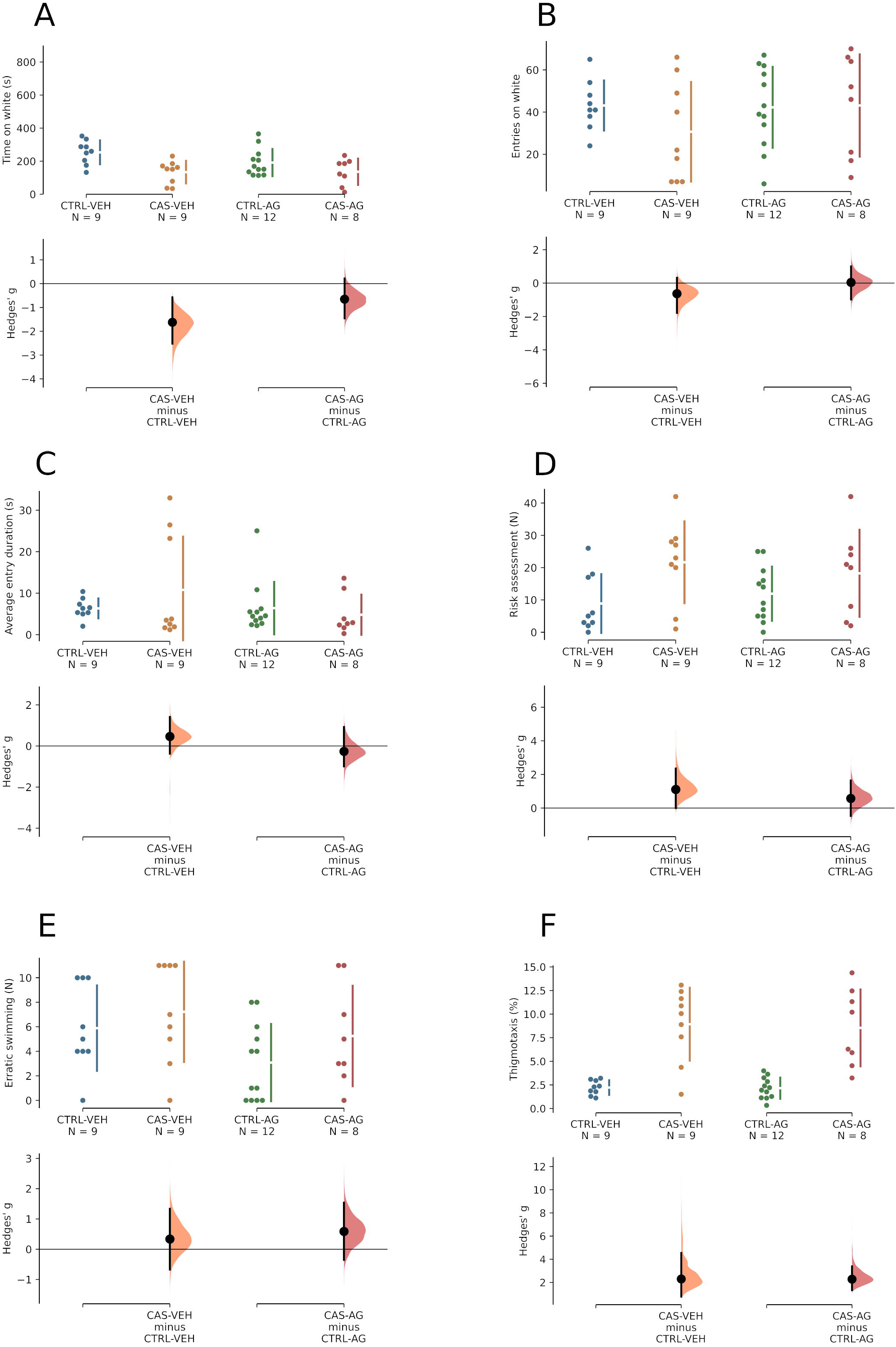
CAS-elicited increases in anxiety-like behavior in the light/dark test 24 h after exposure are not mediated by NOS-2 activated 30 minutes after stress. (A) Scototaxis, indexed as time spent in the white compartment. (B) Total locomotion, indexed as total number of entries in the white compartment. (C) Average entry duration. (D) Total number of risk assessment events. (E) Total number of erratic swimming events. (F) Thigmotaxis, indexed by proportion of time in the white compartment spent near the walls. The raw data is plotted on the upper axes; individual points represent individual animals. Each effect size (Cohen’s *d*) is plotted on the lower axes as a bootstrap sampling distribution. 5000 bootstrap samples were taken; the confidence interval is bias-corrected and accelerated. Mean differences are depicted as dots; 95% confidence intervals are indicated by the ends of the vertical error bars.

When animals were treated with AG 90 min. after CAS, a significant effect of treatment (F _[1, 34]_ = 7.17, p = 0.011), but not drug (F_[1, 34]_ = 0.19, p = 0.666), was found for time on white (Figure 6A). Post-hoc analysis found that CAS decreased time on white (p = 0.007, VEH + CTRL vs. VEH+ CAS), an effect that was blocked by AG (p = 0.995, VEH + CAS vs. AG + CAS). No main or interaction effects were observed on entries on white (treatment: F_[1, 34]_ = 0.06, p = 0.808; drug: F_[1, 34]_ = 0.31, p = 0.584; interaction: F_[1, 34]_ = 1.02, p = 0.319; Figure 6B) or average duration of entries (treatment: F_[1, 34]_ = 2.05, p = 0.161; drug: F_[1, 34]_ = 0.97, p = 0.33; interaction: F_[1, 34]_ = 0.25, p = 0.62; Figure 6C). No main or interaction effects were observed on risk assessment (treatment: F_[1, 34]_ = 0.18, p = 0.671; drug: F_[1, 34]_ = 1.07, p = 0.309; interaction: F_[1, 34]_ = 0.15, p = 0.607; Figure 6D). No main or interaction effects were observed on erratic swimming (treatment: F_[1, 34]_ = 1.09, p = 0.304; drug: F_[1, 34]_ = 0.28, p = 0.603; interaction: F_[1, 34]_ = 0.07, p = 0.797; Figure 6E). Main effects of treatment (F_[1, 34]_ = 13.59, p < 0.001) and drug (F_[1, 34]_ = 14.57, p < 0.001), as well as an interaction effect (F_[1, 34]_ = 12.36, p = 0.001), were found for thigmotaxis (Figure 6F). Post-hoc analysis found that CAS increased thigmotaxis (p < 0.001, VEH + CTRL vs. VEH + CAS), an effect that was blocked by AG (p = 0.999, VEH + CAS vs. AG + CAS).

**Figure 6.**
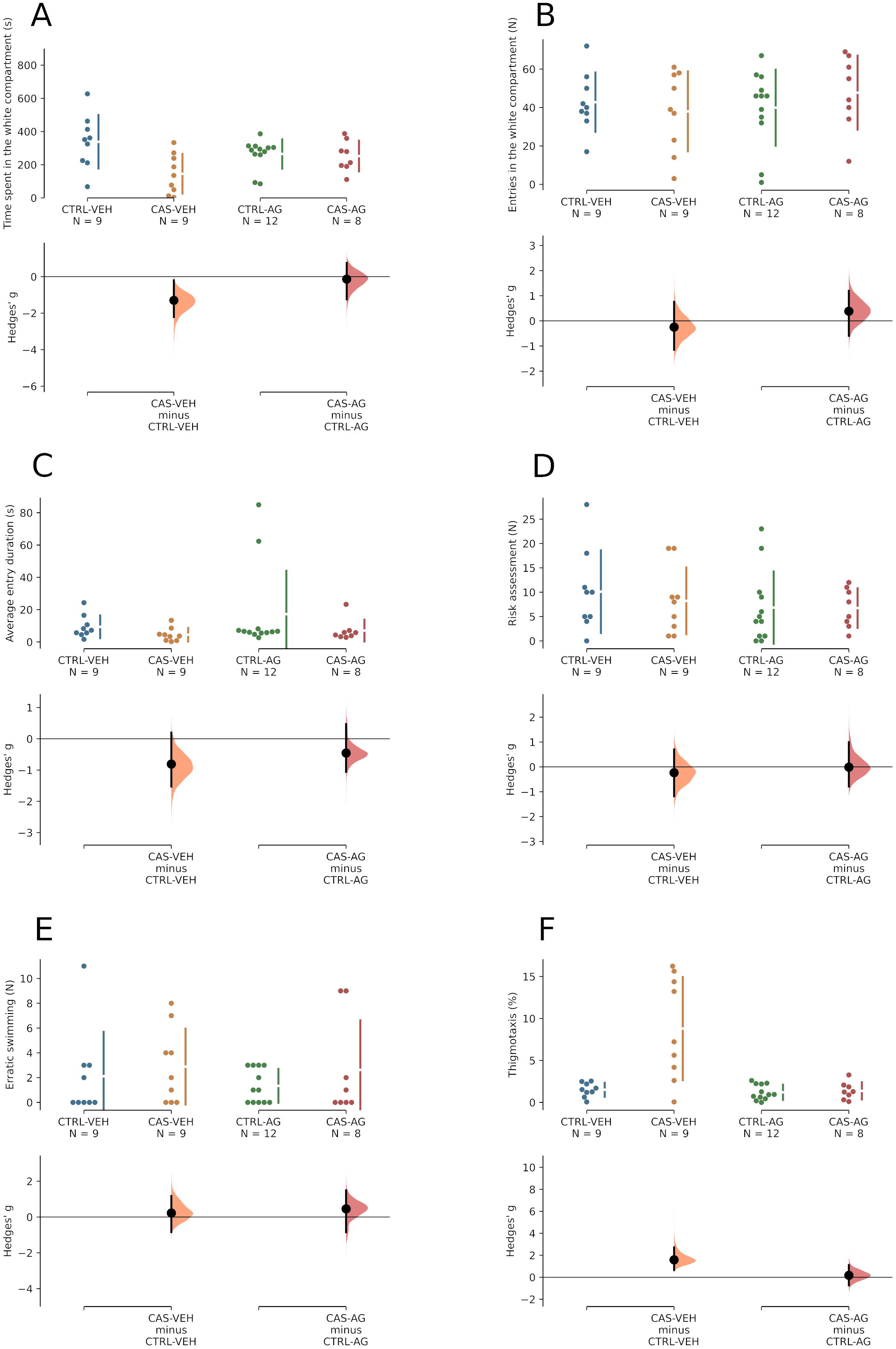
CAS-elicited increases in anxiety-like behavior in the light/dark test 24 h after exposure are mediated by NOS-2 activated 90 minutes after stress. (A) Scototaxis, indexed as time spent in the white compartment. (B) Total locomotion, indexed as total number of entries in the white compartment. (C) Average entry duration. (D) Total number of risk assessment events. (E) Total number of erratic swimming events. (F) Thigmotaxis, indexed by proportion of time in the white compartment spent near the walls. The raw data is plotted on the upper axes; individual points represent individual animals. Each effect size (Cohen’s *d*) is plotted on the lower axes as a bootstrap sampling distribution. 5000 bootstrap samples were taken; the confidence interval is bias-corrected and accelerated. Mean differences are depicted as dots; 95% confidence intervals are indicated by the ends of the vertical error bars.

#### 3.2.3. KCNN channels participate in the second time window of behavioral sensitization: 90 min. after CAS exposure

One of the sources of NOS-2 induction in the central nervous system is the activation of microglial calcium-activated potassium channels (KCNN channels) (Kaushal et al., 2007). To understand the role of KCNN channels in TDS, animals were treated with triarylmethane-34 (TRAM-34), a KCNN blocker, 30 min. or 90 min. after CAS, and their behavior in the LDT was observed 24 h after stress. When animals were treated with TRAM-34 30 min. after stress, a main effect of treatment (F_[1, 41]_ = 9.98, p = 0.003) was found for time on white, but not main effect of drug (F_[1, 41]_ = 0.2, p = 0.661) nor an interaction effect (F_[1, 41]_ = 2.73, p = 0.106) were found for time on white. CAS significantly decreased time on white 24 h after exposure, regardless of drug dose (Figure 7A). No main effects of treatment (F_[1, 41]_ = 0.61, p = 0.438) or drug (F_[1, 41]_ = 0.02, p = 0.893) were found for entries on the white compartment, an an interaction effect was also absent (F_[1, 41]_ = 2.37, p = 0.131; Figure 7B). The same was found for the average duration of entries (treatment: F_[1, 41]_ = 0.3, p = 0.587; drug: F_[1, 41]_ = 0.03, p = 0.863; interaction: F_[1, 41]_ = 3.74, p = 0.06; Figure 7C). A main effect of treatment (F_[1, 41]_ = 28.76, p < 0.001) was found for risk assessment, but a main effect of drug was absent (F_[1, 41]_ = 1.29, p = 0.264); an interaction effect was also absent (F_[1, 41]_ = 0.65, p = 0.423). CAS increased risk assessment 24 h after exposure, regardless of drug dose (Figure 7D). No main nor interaction effects were found for erratic swimming (treatment: F_[1, 41]_ = 0.03, p = 0.867; drug: F_[1, 41]_ = 2.3, p = 0.137; interaction: F_[1, 41]_ = 3.18, p = 0.082; Figure 7E). A main effect of treatment (F_[1, 41]_ = 56.83, p < 0.001), but not drug (F_[1, 41]_ = 0.61, p = 0.44), was found for thigmotaxis; no interaction effect was found (F_[1, 41]_ = 0.16, p = 0.69). In general, CAS increased thigmotaxis 24 h after exposure, regardless of drug dose (Figure 7F). In general, CAS increased anxiety-like behavior 24 h after exposure, and TRAM-34 injected 30 min. after stress did not prevent that effect.

**Figure 7.**
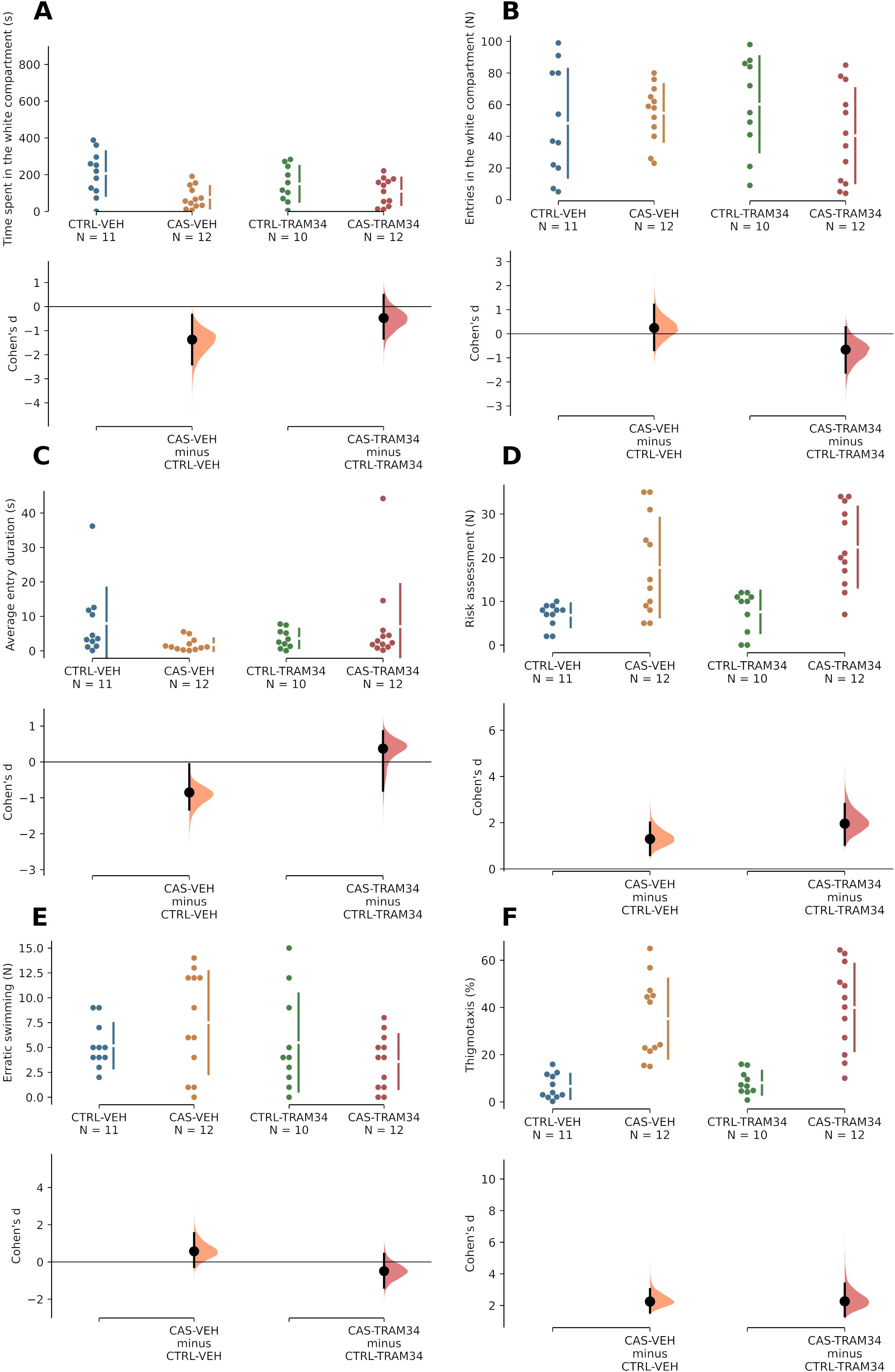
CAS-elicited increases in anxiety-like behavior in the light/dark test 24 h after exposure are not mediated by KCNN channels activated 30 minutes after stress. (A) Scototaxis, indexed as time spent in the white compartment. (B) Total locomotion, indexed as total number of entries in the white compartment. (C) Average entry duration. (D) Total number of risk assessment events. (E) Total number of erratic swimming events. (F) Thigmotaxis, indexed by proportion of time in the white compartment spent near the walls. The raw data is plotted on the upper axes; individual points represent individual animals. Each effect size (Cohen’s *d*) is plotted on the lower axes as a bootstrap sampling distribution. 5000 bootstrap samples were taken; the confidence interval is bias-corrected and accelerated. Mean differences are depicted as dots; 95% confidence intervals are indicated by the ends of the vertical error bars.

When animals were treated with TRAM-34 90 min. after CAS, no main effect of treatment was found for time on white (F_[1, 42]_ = 1.76, p = 0.192), but a main effect of drug (F_[1, 42]_ = 7.8, p = 0.008) and an interaction effect (F_[1, 42]_ = 6.35, p = 0.016) were found (Figure 8A). No main nor interaction effects were found for entries on white (treatment: F_[1, 42]_ = 0.01, p = 0.984; drug: F_[1, 42]_ = 0.3, p = 0.589; interaction: F_[1, 42]_ = 0.19, p = 0.664; Figure 8B) or average entry duration (treatment: F_[1, 42]_ = 0.72, p = 0.402; drug: F_[1, 42]_ = 0.1, p = 0.755; interaction: F_[1, 42]_ = 0.13, p = 0.719; Figure 8C). Main effects of treatment (F_[1, 42]_ = 8.89, p = 0.005) and drug (F_[1, 42]_ = 4.1, p = 0.049) were found for risk assessment (Figure 8D); an interaction effect was also found (F _[1, 42]_ = 4.37, p = 0.043). Post-hoc Tukey’s HSD test found that CAS increased risk assessment (p = 0.004, CTRL + VEH vs. CAS + VEH), an effect that was prevented by TRAM-34 (p = 0.024, CAS + VEH vs. CAS + TRAM-34). No main nor interaction effects were found for erratic swimming (treatment: F[1, 42] = 2.87, p = 0.098; drug: F_[1, 42]_ = 0.02, p = 0.878; interaction: F_[1, 42]_ = 0.3, p = 0.584; Figure 8E). Main effects of treatment (F_[1, 42]_ = 11.77, p = 0.001) and drug (F_[1, 42]_ = 4.31, p = 0.044) were found for thigmotaxis; an interaction effect was also found (F_[1, 42]_ = 5.79, p = 0.021; Figure 8F). Post-hoc analysis found that CAS increased thigmotaxis (p < 0.001, CTRL + VEH vs. CAS + VEH), an effect that was prevented by TRAM-34 (p = 0.012, CAS + VEH vs. CAS + TRAM-34). Thus, in general, TRAM-34, when injected 90 min. after stress, prevented TDS in zebrafish.

**Figure 8.**
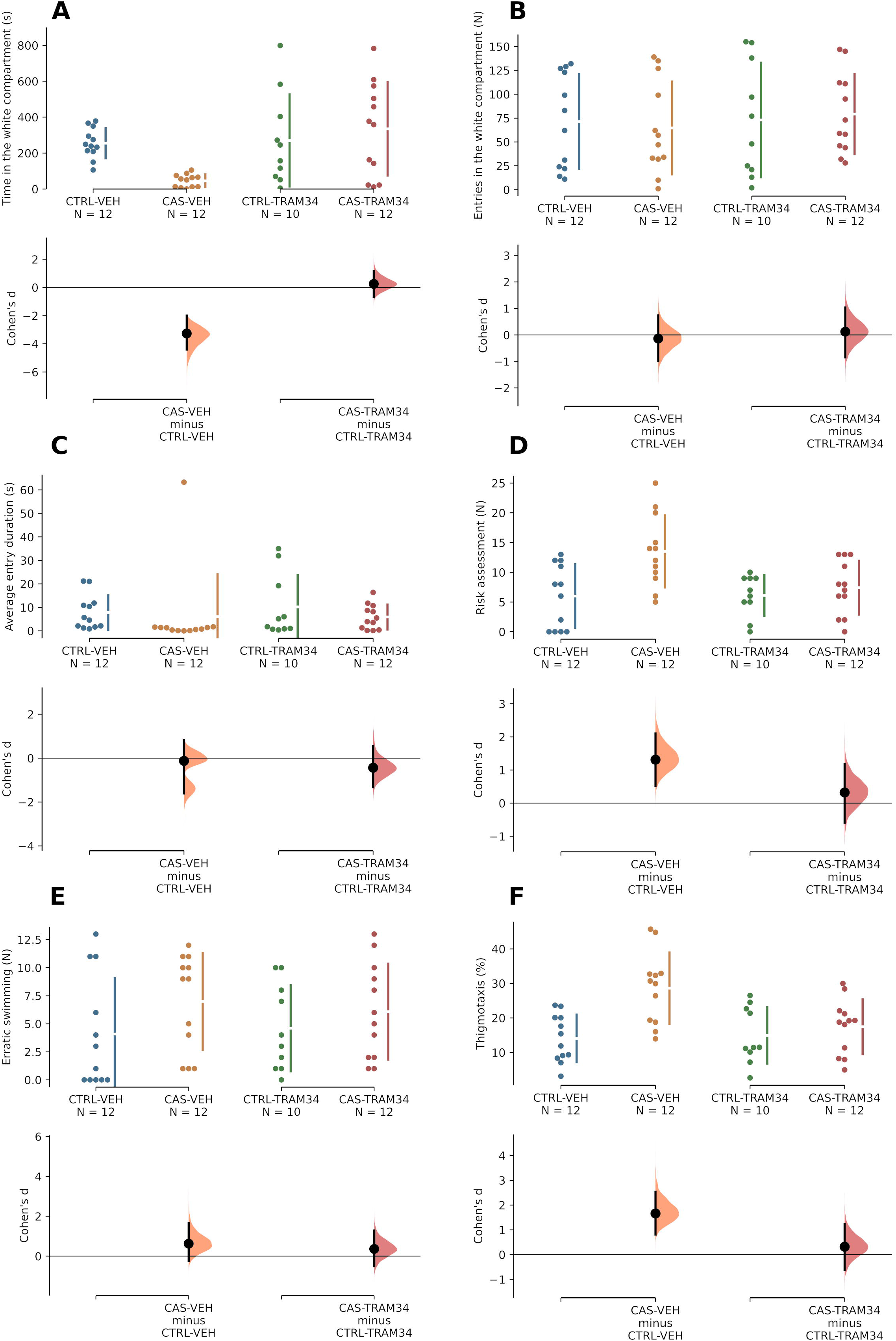
CAS-elicited increases in anxiety-like behavior in the light/dark test 24 h after exposure are mediated by KCNN channels activated 90 minutes after stress. (A) Scototaxis, indexed as time spent in the white compartment. (B) Total locomotion, indexed as total number of entries in the white compartment. (C) Average entry duration. (D) Total number of risk assessment events. (E) Total number of erratic swimming events. (F) Thigmotaxis, indexed by proportion of time in the white compartment spent near the walls. The raw data is plotted on the upper axes; individual points represent individual animals. Each effect size (Cohen’s *d*) is plotted on the lower axes as a bootstrap sampling distribution. 5000 bootstrap samples were taken; the confidence interval is bias-corrected and accelerated. Mean differences are depicted as dots; 95% confidence intervals are indicated by the ends of the vertical error bars.

#### 3.2.4. Participation of guanylate cyclase on sensitization in the second time window of behavioral sensitization

The “canonical” downstream effectors of NO are dependent on cyclic guanosine monophosphate (cGMP). Telencephalic cGMP is involved in memory consolidation in avoidance conditioning in goldfish (Xu et al., 2009), suggesting a role for this second messenger in processes that are relevant to PTSD. In order to assess the role of guanylate cyclase (GC), the NO-dependent enzyme responsible for converting GMP into cGMP, on behavioral sensitization during the incubation period, animals were treated with ODQ, a specific GC blocker, 30 min and 90 min. after CAS, and their behavior in the LDT was observed 24 h after stress. When animals were treated with ODQ 30 min. after stress, no main effects of treatment on time on white were observed (F _[1,89]_ = 0.37, p = 0.544), but a significant effect of drug (F_[1, 89]_ = 9.49, p = 0.003) and an interaction effect (F_[1, 89]_ = 4.37, p = 0.039) were found (Figure 9A). Post-hoc analysis found that CAS decreased time on white (p = 0.046, VEH + CTRL vs. VEH + CAS), an effect that was blocked by ODQ (p = 0.003, VEH + CAS vs. ODQ + CAS). No main effects or interaction effects were found for entries in the white compartment (treatment: F_[1, 89]_ = 0.06, p = 0.806; drug: F_[1, 89]_ = 0.04, p = 0.834; interaction: F_[1, 89]_ = 0.2, p = 0.654; Figure 9B). No main effects or interaction effects were found for the average duration of entries in the white compartment (treatment: F_[1, 89]_ = 0.02, p = 0.891; drug: F_[1, 89]_ = 0.19, p = 0.661; interaction: F_[1, 89]_ = 3.25, p = 0.075; Figure 9C). No main effects or interaction effects were found for risk assessment (treatment: F_[1, 89]_ = 0.09, p = 0.764; drug: F_[1, 89]_ = 0.02, p = 0.897; interaction: F_[1, 89]_ = 1.24, p = 0.269; Figure 9D). A main effect of treatment (F[1, 89] = 10.15, p = 0.002), but not drug (F[1, 89] = 3.02, p = 0.086), was found for erratic swimming (Figure 9E); an interaction effect was also found (F_[1, 89]_ = 4.73, p = 0.032). Post-hoc analysis found that CAS increased erratic swimming (p = 0.001, VEH + CTRL vs. VEH + CAS), an effect that was partially blocked by ODQ (p = 0.038, VEH + CAS vs. ODQ + CAS). No main or interaction effects were found for thigmotaxis (treatment: F_[1, 89]_ = 0.26, p = 0.612; drug: F_[1, 89]_ = 0, p = 0.969; interaction: F_[1, 89]_ = 3.84, p = 0.053; Figure 9F).

**Figure 9.**
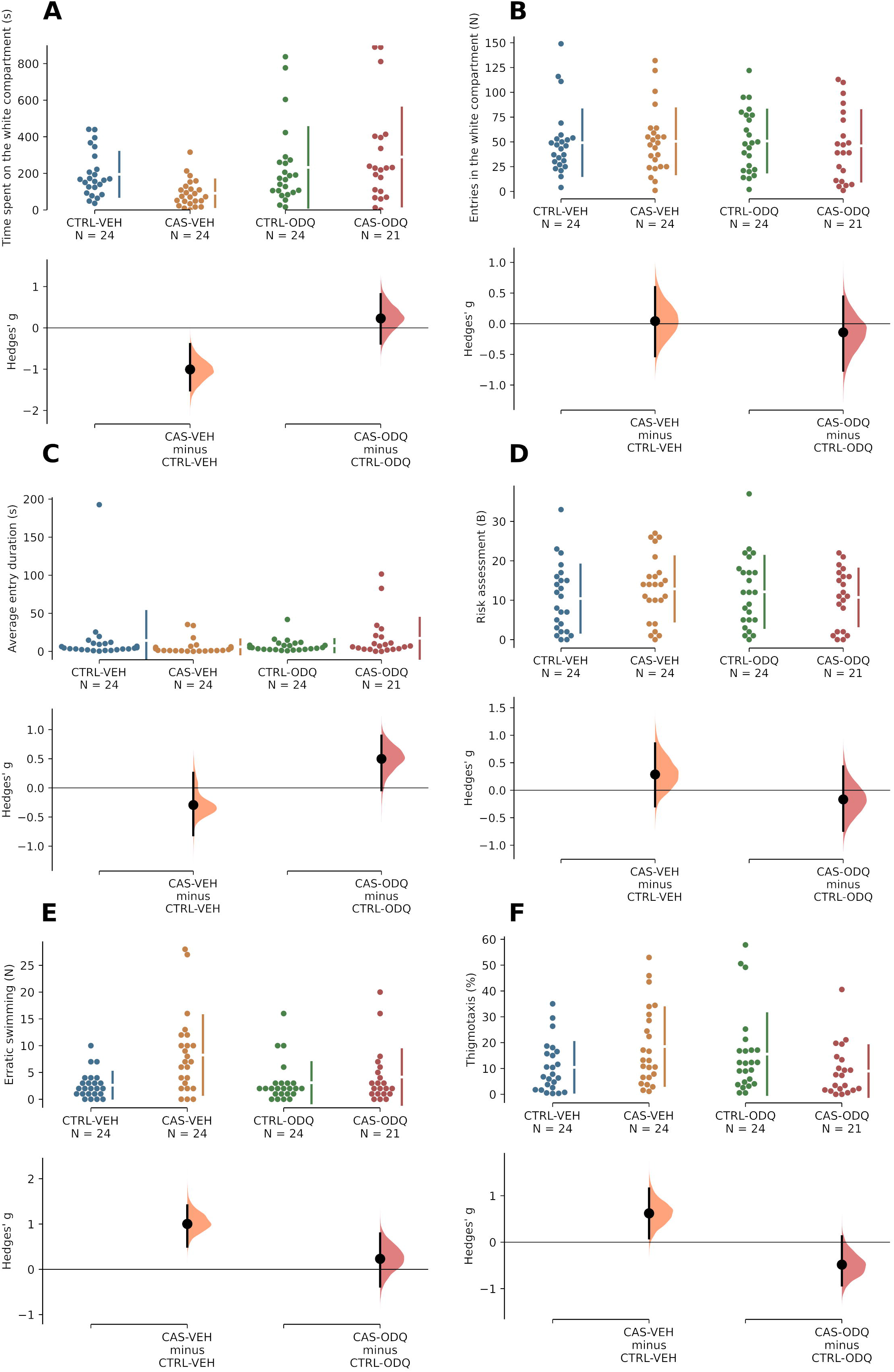
CAS-elicited increases in anxiety-like behavior in the light/dark test 24 h after exposure are not mediated by guanylate cyclase activated 30 minutes after stress. (A) Scototaxis, indexed as time spent in the white compartment. (B) Total locomotion, indexed as total number of entries in the white compartment. (C) Average entry duration. (D) Total number of risk assessment events. (E) Total number of erratic swimming events. (F) Thigmotaxis, indexed by proportion of time in the white compartment spent near the walls. The raw data is plotted on the upper axes; individual points represent individual animals. Each effect size (Cohen’s *d*) is plotted on the lower axes as a bootstrap sampling distribution. 5000 bootstrap samples were taken; the confidence interval is bias-corrected and accelerated. Mean differences are depicted as dots; 95% confidence intervals are indicated by the ends of the vertical error bars.

When animals were treated with ODQ 90 min. after stress, no main effects of treatment (F_[1, 78]_ = 1.94, p = 0.167) were found for time on white (Figure 10A), but a main effect of drug was found (F_[1, 78]_ = 5.4, p = 0.023); an interaction effect was also found (F_[1, 78]_ = 4.6, p = 0.035). Post-hoc analysis found that CAS decreased time on white (p = 0.047, VEH + CTRL vs. VEH + CAS), an effect that was blocked by ODQ (p = 0.009, VEH + CAS vs. ODQ + CAS). No main or interaction effects were found for entries on white (treatment: F_[1, 78]_ = 0.02, p = 0.901; drug: F_[1 78]_ = 0.12, p = 0.735; interaction: F_[1, 78]_ = 0.01, p = 0.988; Figure 10B) or average entry duration (treatment: F_[1, 78]_ = 0.49, p = 0.487; drug: F_[1 78]_ = 0.15, p = 0.7; interaction: F_[1, 78]_ = 1.93, p = 0.169; Figure 10C). A main effect of treatment was found for risk assessment (F_[1, 78]_ = 5.28, p = 0.024); a main effect of drug was found for this variable as well (F_[1, 78]_ = 9.46, p = 0.003), as well as an interaction effect (F_[1, 78]_ = 4.05, p = 0.048). ODQ decreased risk assessment by itself (p = 0.003), and CAS increased risk assessment (p = 0.019, VEH + CTRL vs. VEH + CAS), an effect that was blocked by ODQ (p = 0.002, VEH + CAS vs. ODQ + CAS) (Figure 10D). No main effect of treatment was found for erratic swimming (F_[1, 78]_ = 3.17, p = 0.079), but a main effect of drug was present (F_[1, 78]_ = 4.18, p = 0.044); ODQ increased erratic swimming (p = 0.044; Figure 10E). Finally, main effects of treatment (F_[1, 78]_ = 18.66, p < 0.001) and drug (F_[1, 78]_ = 6.65, p = 0.012) were found for thigmotaxis, and an interaction effect was also present (F_[1, 78]_ = 14, p < 0.001). CAS increased thigmotaxis (p < 0.001, VEH + CTRL vs. VEH + CAS), an effect that was blocked by ODQ (p < 0.001, VEH + CAS vs. ODQ + CAS).

**Figure 10.**
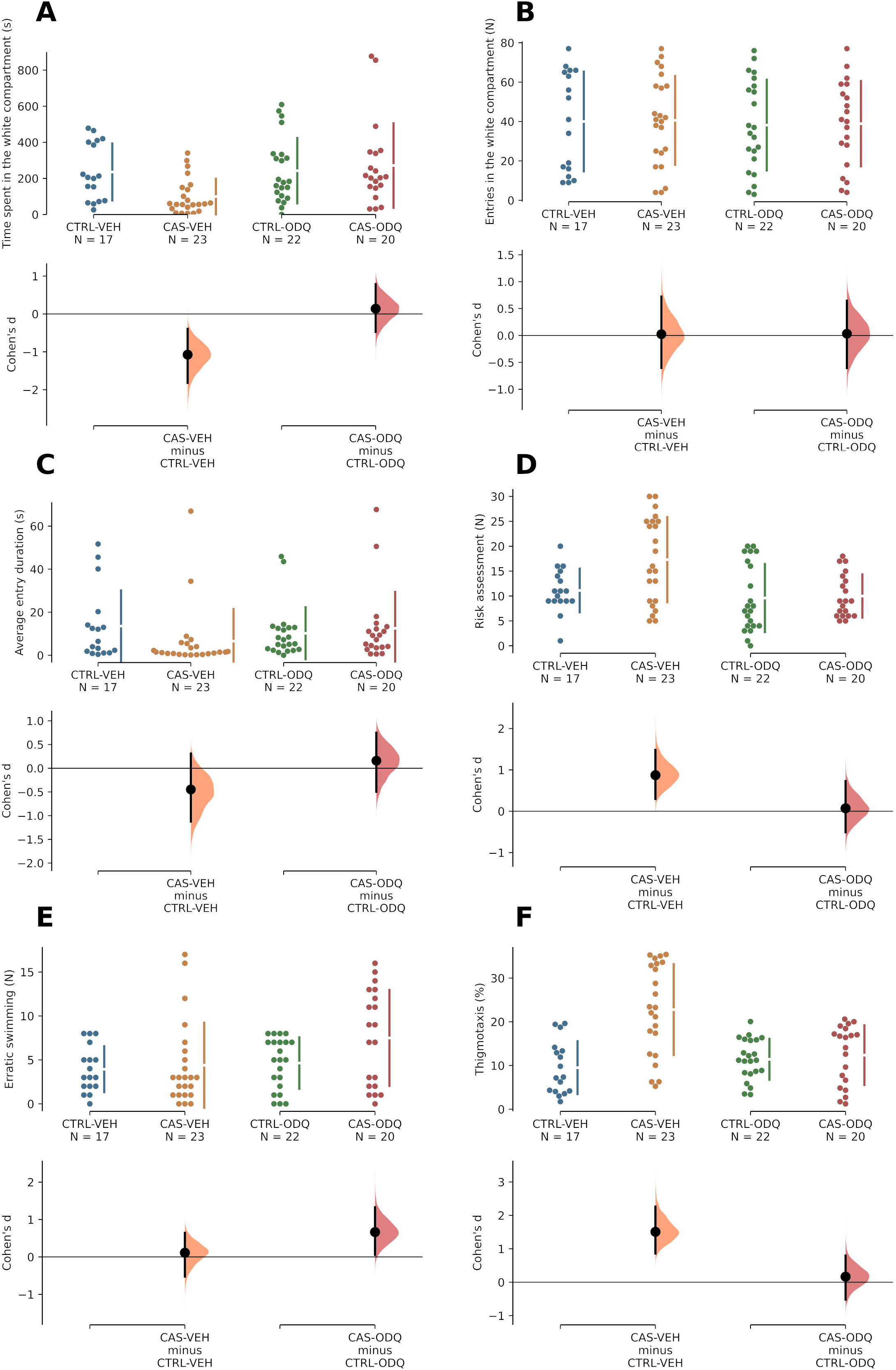
CAS-elicited increases in anxiety-like behavior in the light/dark test 24 h after exposure are mediated by guanylate cyclase activated 90 minutes after stress. (A) Scototaxis, indexed as time spent in the white compartment. (B) Total locomotion, indexed as total number of entries in the white compartment. (C) Average entry duration. (D) Total number of risk assessment events. (E) Total number of erratic swimming events. (F) Thigmotaxis, indexed by proportion of time in the white compartment spent near the walls. The raw data is plotted on the upper axes; individual points represent individual animals. Each effect size (Cohen’s *d*) is plotted on the lower axes as a bootstrap sampling distribution. 5000 bootstrap samples were taken; the confidence interval is bias-corrected and accelerated. Mean differences are depicted as dots; 95% confidence intervals are indicated by the ends of the vertical error bars.

#### 3.2.5. Participation of cGMP-dependent channels on sensitization in the second time window of behavioral sensitization

Hyperpolarization-activated cyclic nucleotide-gated (HCN) channels are important downstream targets of cGMP, and have been shown to be important in synaptic plasticity. To understand the participation of these channels in TDS, animals were treated with ivabradine (IVA), a non-selective HCN channel blocker, 30 min or 90 min after CAS. When animals were treated 30 min after CAS, a significant main effect of treatment (F_[1, 40]_ = 4.35, p = 0.044) was found for time on white, but no main effect of drug was found (F_[1, 40]_ = 1.23, p = 0.275); an interaction effect was also absent (F_[1, 40]_ = 0.07, p = 0.795). Thus, CAS decreased time on white in both vehicle- and ivabradine-treated animals (Figure 11A). No main or interaction effects were found for number of entries (treatment: F_[1, 40]_ = 0.26, p = 0.611; drug: F_[1, 40]_ = 1.83, p = 0.184; interaction: F_[1, 40]_ = 1; p = 0.323; Figure 11B) or average entry duration (treatment: F_[1, 40]_ = 1.13, p = 0.293; drug: F_[1, 40]_ = 1.02, p = 0.319; interaction: F_[1, 40]_ = 0.81, p = 0.374; Figure 11C). No main effect of treatment was found for risk assessment (F_[1, 40]_ = 0.05, p = 0.825), but a significant main effect of drug was found for this endpoint (F_[1, 40]_ = 7.12, p = 0.011); an interaction effect was absent (F_[1, 40]_ = 0.79. p = 0.379). In general, ivabradine-treated animals showed higher risk assessment, regardless of whether exposed or not to CAS (Figure 11D). No main or interaction effects were found for erratic swimming (treatment: F_[1, 40]_ = 0.06, p = 0.801; drug: F_[1, 40]_ = 0.35, p = 0.558; interaction: F_[1, 40]_ = 1.83; p = 0.184; Figure 11E). No main effects of treatment (F_[1, 40]_ = 2.14, p = 0.152) were found for thigmotaxis, but a significant main effect of drug was found (F_[1, 40]_ = 4.18, p = 0.048); a significant interaction effect was also found (F_[1, 40]_ = 9.48, p = 0.004; Figure 11F). Post-hoc analysis found that CAS increased thigmotaxis (p = 0.013, CTRL + VEH vs. CAS + VEH), an effect that was prevented by ivabradine (p = 0.004, CAS + VEH vs. CAS + IVA; Figure 11F). Thus, HCN channels appear not to be involved in TDS in the first time window, although a sensitizing effect on thigmotaxis was observed.

**Figure 11.**
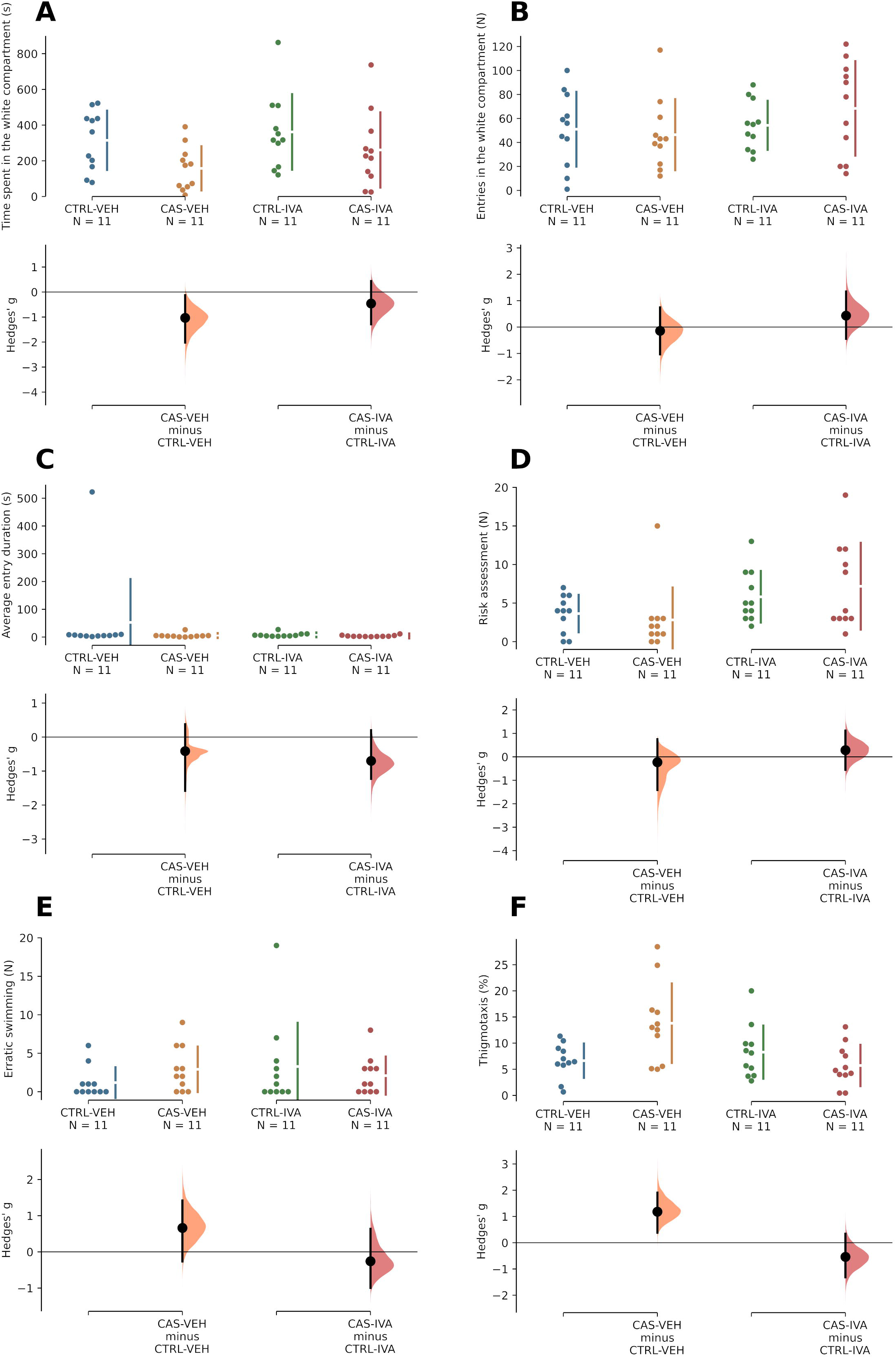
CAS-elicited increases in anxiety-like behavior in the light/dark test 24 h after exposure are not mediated by HCN channels activated 30 minutes after stress. (A) Scototaxis, indexed as time spent in the white compartment. (B) Total locomotion, indexed as total number of entries in the white compartment. (C) Average entry duration. (D) Total number of risk assessment events. (E) Total number of erratic swimming events. (F) Thigmotaxis, indexed by proportion of time in the white compartment spent near the walls. The raw data is plotted on the upper axes; individual points represent individual animals. Each effect size (Cohen’s *d*) is plotted on the lower axes as a bootstrap sampling distribution. 5000 bootstrap samples were taken; the confidence interval is bias-corrected and accelerated. Mean differences are depicted as dots; 95% confidence intervals are indicated by the ends of the vertical error bars.

When animals were treated with ivabradine 90 min after CAS, main effects of treatment (F[1, 39] = 6.44, p = 0.015) and drug (F[1, 39] = 9.6, p = 0.004) were found for time on white (Figure 12A); an interaction effect was also found (F[1, 39] = 4.67, p = 0.037). Post-hoc analysis found that CAS decreased time on white (p = 0.007, CTRL + VEH vs. CAS + VEH), an effect that was prevented by ivabradine (p = 0.005, CAS + VEH vs. CAS + IVA). No main or interaction effects were found for entries on white (treatment: F[1, 39] = 1.59, p = 0.215; drug: F[1, 39] = 3.77, p = 0.059; interaction: F[1, 39] = 0.25, p = 0.619; Figure 12B). No main effects of treatment were found for average entry duration (F[1, 39] = 0.1, p = 0.75), but a main effect of drug was found for this variable (F[1, 39] = 25.1, p < 0.001); since an interaction effect was not found (F[1, 39] = 3.33, p = 0.075), we conclude that ivabradine increased average entry duration across treatment conditions (Figure 12C). No main or interaction effects were found for risk assessment (treatment: F[1, 39] = 1.48, p = 0.23; drug: F[1, 39] = 0.01, p = 0.985; interaction: F[1, 39] = 0.82, p = 0.37; Figure 12E), thigmotaxis (treatment: F[1, 39] = 0.22, p = 0.645; drug: F[1, 39] = 2.49, p = 0.122; interaction: F[1, 39] = 0.38, p = 0.541; Figure 12E), erratic swimming (treatment: F[1, 39] = 0.42, p = 0.52; drug: F[1, 39] = 3.09, p = 0.087; interaction: F[1, 39] = 1.69, p = 0.201; Figure 12F), or freezing (treatment: F[1, 39] = 4 p = 0.052; drug: F[1, 39] = 0.06, p = 0.813; interaction: F[1, 39] = 0.26, p = 0.61; Figure 12G). Thus, HCN channels appear to be involved in TDS in the second time window.

**Figure 12.**
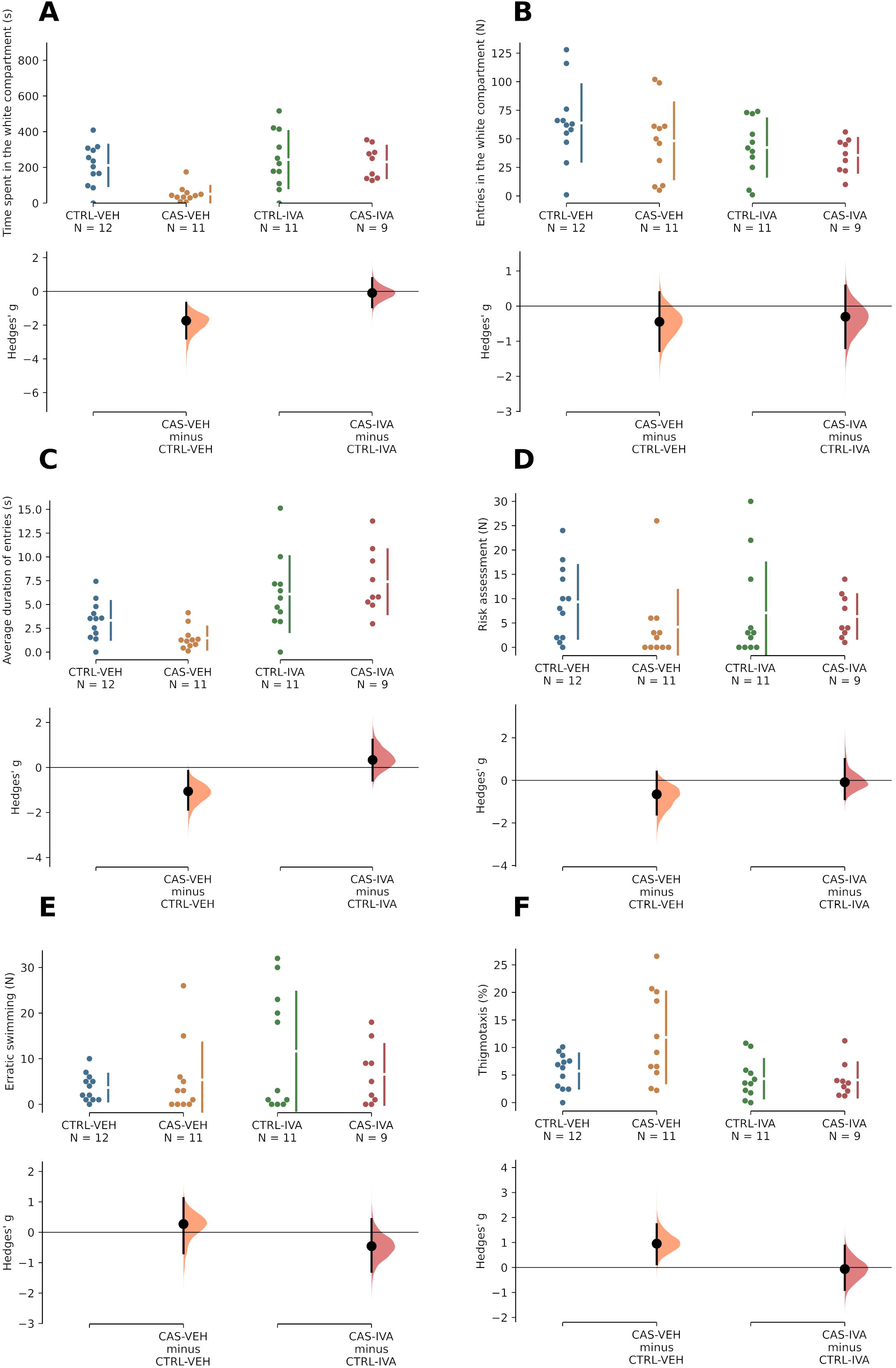
CAS-elicited increases in anxiety-like behavior in the light/dark test 24 h after exposure are mediated by HCN channels activated 90 minutes after stress. (A) Scototaxis, indexed as time spent in the white compartment. (B) Total locomotion, indexed as total number of entries in the white compartment. (C) Average entry duration. (D) Total number of risk assessment events. (E) Total number of erratic swimming events. (F) Thigmotaxis, indexed by proportion of time in the white compartment spent near the walls. The raw data is plotted on the upper axes; individual points represent individual animals. Each effect size (Cohen’s *d*) is plotted on the lower axes as a bootstrap sampling distribution. 5000 bootstrap samples were taken; the confidence interval is bias-corrected and accelerated. Mean differences are depicted as dots; 95% confidence intervals are indicated by the ends of the vertical error bars.

## 4. Discussion

The present paper described the roles of molecules in the NO pathway on behavioral sensitization in time-dependent sensitization (TDS), a model for post-traumatic stress disorder (PTSD), in zebrafish. Following on previous results that suggest two time windows of NO-dependent sensitization in the “incubation” period of TDS, we show that acute exposure to conspecific alarm substance (CAS) elicits behavioral sensitization 24 h after stress; increased extracellular levels of glutamate in the telencephalon immediately after stress and 30 min. after stress; increased nitrite (NO_X_^-^) levels in the telencephalon immediately after exposure, 30 min. after exposure, 90 min. after exposure, and 24 h after exposure; and increased NO_X_^-^ levels in the head kidney 30 and 90 min. after stress. We also show that the increases in telencephalic NO_X_^-^ levels 90 min. after stress (but not in the first time window) were dependent on NOS-2, while increases in the head kidney were not. Using drugs to block different levels of the pathway, we showed that TDS is dependent on the activation of KCNN channels in the second time window (90 min. after stress), NOS-1 activation in the first time window (30 min. after stress), NOS-2 activation in the second time window, guanlyate cyclase (GC) activation in both time windows, and cGMP-dependent channel activation in the second time window (Figure 13).

**Figure 13.**
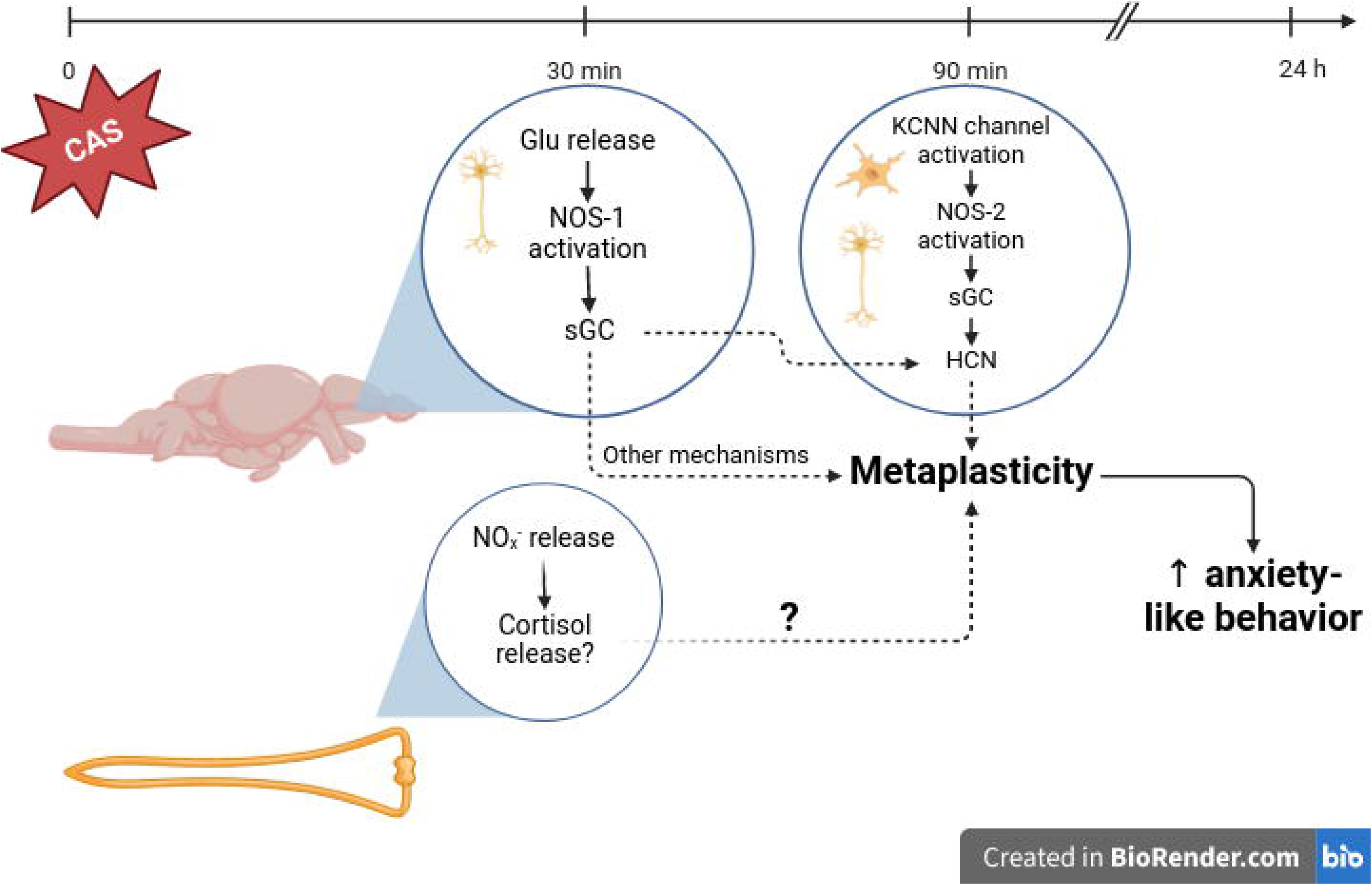
A theoretical model of the participation of NOS isoforms and upstream and downstream mediators in the NO cascade during the consolidation of time-dependent sensitization. Conspecific alarm substance (CAS) elicits glutamate release in the forebrain, which leads to the activation of the NOS-1-sGC pathway in forebrain neurons 30 min. after stress. Simultaneously, nitrergic signalling in the head kidney leads to the release of cortisol and/or catecholamines. After 90 min., the activation of KCNN channels in forebrain neurons and microglia leads to the synthesis and activation of NOS-2. The activation of soluble guanylate cyclase (sGC) by nitric oxide produced by both NOS-1 and NOS-2 leads to the activation of HCN channels, contributing to metaplasticity of limbic circuits. The sensitization of these circuits by metaplasticity will result in increased anxiety-like behavior when animals are tested 24 h after CAS exposure.

Increases in glutamate levels in the telencephalon after stress suggest that one of the pathways involved in TDS is the NDMA-R-NOS-1 pathway. NMDA-R activation has been shown to increase anxiety-like behavior in zebrafish (Herculano et al., 2015), and to be necessary for aversive learning in zebrafish (Blank et al., 2009; Xu et al., 2007b) and goldfish (Xu et al., 2003), processes which can be relevant to TDS. While operations of neurobehavioral sensitization are thought to involve both associative and non-associative processes (Daskalakis et al., 2013b), the activation of NMDA-Rs is likely to be involved in TDS, leading to the activation of downstream effectors, such as NOS-1. Indeed, in a model of behavioral sensitization after predator exposure in rats, blocking NMDA-Rs before predator exposure, but not after exposure, blocks the sensitization (Blundell et al., 2005; Blundell & Adamec, 2007). Importantly, in the present work, blocking NOS-1 30 min. after stress inhibited TDS, suggesting that, in the first moments of the incubation period, the NDMA-R-NOS-1 pathway is necessary for TDS.

The increases in glutamate are followed by increases in nitrite-levels in the telencephalon, an increase that was sustained up to 24 h after stress. Following on previous work which showed that blocking NOS isoforms in two time windows during the incubation period (30 min. and 90 min. after CAS), we tested the role of NOS-2 on these increases, showing that this isoform participates in the second time window. In a rodent model of TDS, hippocampal NOS activity was increased 21 days after stress, an effect that was blocked by blocking NOS-2 (Harvey et al., 2004). We also found that blocking NOS-2 90 min after stress prevents TDS, corroborating the idea that NOS-2-dependent increases in telencephalic NO production are related to behavioral sensitization in PTSD-relevant models. NOS-2 has previously been shown to participate in the increases in anxiety-like behavior in zebrafish and rats exposed to ethanol withdrawal (Bonassoli et al., 2011; da Silva Chaves et al., 2020), suggesting that this inducible isoform also participates in other long-term sensitization processes related to defensive behavior. Since we also showed that CAS leads to increases in nitrite levels in the head kidney, in which cortisol-producing interrenal cells are located, we tested whether NOS-2 participated in these processes, and found no evidence for this hypothesis; as a result, the role of NOS-2 in the second time window is more likely to be directly related to the limbic telencephalon, and not to its indirect modulation by e.g. cortisol secretion. The role of interrenal nitric oxide in neuroendocrine effects of TDS are not yet known.

While the role of the NMDA receptor in the recruitment of NOS-1 is well-established, the upstream inducers of NOS-2 are diverse (Cinelli et al., 2020). The major pathways involve either the activation of NF-κB or the JAK-STAT pathway by cytokines. In microglia, the activation of KCNN channels are an important mediator of microglia activation and NOS-2 activation (Kaushal et al., 2007). When animals were treated with the KCNN blocker TRAM-34 90 in after stress, TDS was prevented, implicating these channels in the long-term behavioral effects of CAS. While the exact cellular location of the effects of TRAM-34 in the present experiments are unknown, we suggest that stress activates first the NMDA-NOS-1 pathway, and later a KCNN-NOS-2 pathway, to induce sustained production of NO in the limbic telencephalon, sensitizing circuits of the aversive brain circuit and increasing anxiety-like behavior (Figure 13). Supporting the idea that this pathway is at least partially related to microglial activity, a study on zebrafish found that a combined 90-min stress that leads to TDS also led to upregulation of the expression of microglial markers as pro-inflammatory cytokines (Yang et al., 2020).

In addition to determining upstream mechanisms and NOS isoforms involved in TDS, we also sought to determine downstream effector pathways. Blocking GC prevented TDS in both time windows, implicating this enzyme, and subsequent cGMP signaling, as downstream effectors of both NOS-1- and NOS-2-dependent behavioral sensitization mechanisms. We also found that blocking nucleotide-dependent channels, which can be activated by cAMP or cGMP, blocked TDS in the second time window, implicating these channels in the latter moment of incubation.

In general, our results show the participation of multiple NO-related pathways in different time windows of the early incubation period of TDS. We suggest that CAS first leads to the release of glutamate and subsequent activation of the NMDA-R-NOS-1 pathway in the limbic telencephalon, a process which, via sGC activation and cGMP synthesis, initiates metaplasticity (Figure 13). Simultaneously, activation of a NO pathway in the head kidney can lead to the release of either cortisol or catecholamines, which can synergistically contribute with metaplasticity. In a second wave of nitrergic activity, the activation of KCNN channels in microglia leads to NOS-2 synthesis and activation. Both the cGMP synthesized in the NMDA-R-NOS-1 pathway and the KCNN-NOS-2 pathway lead to the activation of HCN channels, which ultimately contribute to metaplasticity. Moreover, sustained NO production (up to 24 h after stress), mediated by NOS-2, can also lead to other mechanisms, including nitrosative stress, which sensitize limbic circuits; the behavioral result is increased anxiety-like behavior (Figure 13).

The present work lends support to a model in which both NMDA-R-NOS-1 and KCNN-NOS-2 pathways participate in the incubation of time-dependent sensitization, a behavioral model which is relevant to understand trauma- and stressor-related disorders. In combination with previous results demonstrating the delayed behavioral effects of stressors (i.e., TDS) in zebrafish (Lima et al., 2016; Maximino et al., 2018; Theron et al., 2023; Yang et al., 2020), the role of NOS isoforms in the incubation of TDS (Lima et al., 2015), and the role of NO pathways in learning mechanisms of the limbic telencephalon of teleost fish (Xu et al., 2003, 2009), the present results reinforce the hypothesis that nitrergic modulation of (meta)plasticity in the limbic telencephalon is a potential mechanism in the development of trauma- and stressor-related disorders. Further work is needed to understand the electrophysiological bases of these mechanisms, as well as to test the translatability of this model for patients.

## Acknowledgments

Authors are thankful to Dr. Valney Mara Gomes Conde for the generous donation of reagents. Funding for this research was possible due to a Conselho Nacional de Desenvolvimento Científico e Tecnológico (CNPq/Brazil) grant to MLM (grant # 423735/2016-0).

## References

Abraham, W. C. (2008). Metaplasticity: Tuning synapses and networks for plasticity. Nature Reviews Neuroscience, 9(5), 387–387. 10.1038/nrn2356

Abraham, W. C., & Richter-Levin, G. (2018). From Synaptic Metaplasticity to Behavioral Metaplasticity. Neurobiology of Learning and Memory, 154, 1–4. 10.1016/j.nlm.2018.08.015

Abraham, W. C., & Tate, W. P. (1997). Metaplasticity: A new vista across the field of synaptic plasticity. Progress in Neurobiology, 52(4), 303–323. 10.1016/S0301-0082(97)00018-X

Adamec, R. E., Blundell, J., & Burton, P. (2005a). Neural circuit changes mediating lasting brain and behavioral response to predator stress. Neuroscience & Biobehavioral Reviews, 29, 1225–1241. 10.1016/j.neubiorev.2005.05.007

Adamec, R. E., Blundell, J., & Burton, P. (2005b). Role of NMDA receptors in the lateralized potentiation of amygdala afferent and efferent neural transmission produced by predator stress. Physiology & Behavior, 86(1), 75–91. 10.1016/j.physbeh.2005.06.026

Adamec, R. E., Blundell, J., & Collins, A. (2001). Neural plasticity and stress induced changes in defense in the rat. Neuroscience & Biobehavioral Reviews, 25, 721–744. 10.1016/S0149-7634(01)00053-7

Adamec, R. E., Blundell, J., Strasser, K., & Burton, P. (2006). Mechanisms of Lasting Change in Anxiety Induced by Severe Stress. Em N. Kato, M. Kawata, & R. K. Pitman (Orgs.), PTSD (p. 61–81). Springer Japan. 10.1007/4-431-29567-4_7

Adamec, R. E., Kent, P., Anisman, H., Shallow, T., & Merali, Z. (1998). Neural plasticity, neuropeptides and anxiety in animals – Implications for understanding and treating affective disorder following traumatic stress in humans. Neuroscience & Biobehavioral Reviews, 23, 301–318.

Akar, F. Y., Ulak, G., Tanyeri, P., Erden, F., Utkan, T., & Gacar, N. (2007). 7-Nitroindazole, a neuronal nitric oxide synthase inhibitor, impairs passive-avoidance and elevated plus-maze memory performance in rats. *Pharmacology*, Biochemistry and Behavior, 87, 434–443. 10.1016/j.pbb.2007.05.019

Bacila, I., Cunliffe, V. T., & Krone, N. P. (2021). Interrenal development and function in zebrafish. Molecular and Cellular Endocrinology, 535, 111372. 10.1016/j.mce.2021.111372

Baker, D. G., Nievergelt, C. M., & O’Connor, D. T. (2012). Biomarkers of PTSD: Neuropeptides and immune signaling. Neuropharmacology, 62(2), 663–673. 10.1016/j.neuropharm.2011.02.027

Blank, M., Guerim, L. D., Cordeiro, R. F., & Vianna, M. R. M. (2009). A one-trial inhibitory avoidance task to zebrafish: Rapid acquisition of an NMDA-dependent long-term memory. Neurobiology of Learning and Memory, 92, 529–534. 10.1016/j.nlm.2009.07.001

Blundell, J., & Adamec, R. E. (2007). The NMDA receptor antagonist CPP blocks the effects of predator stress on pCREB in brain regions involved in fearful and anxious behavior. Brain Research, 36, 59–76. 10.1016/j.brainres.2006.09.078

Blundell, J., Adamec, R. E., & Burton, P. (2005). Role of NMDA receptors in the syndrome of behavioral changes produced by predator stress. Physiology & Behavior, 86, 233–243. 10.1016/j.physbeh.2005.07.012

Bonassoli, V. T., Milani, H., & Oliveira, R. M. W. de. (2011). Ethanol withdrawal activates nitric oxide-producing neurons in anxiety-related brain areas. Alcohol, 45, 641–652. 10.1016/j.alcohol.2010.11.007

Bruenig, D., Morris, C. P., Mehta, D., Harvey, W., Lawford, B., Young, R. M., & Voisey, J. (2017). Nitric oxide pathway genes (NOS1AP and NOS1) are involved in PTSD severity, depression, anxiety, stress and resilience. Gene, 625, 42–48. 10.1016/j.gene.2017.04.048

Calabrese, V., Mancuso, C., Calvani, M., Rizzarelli, E., Butterfield, D. A., & Stella, A. M. G. (2007). Nitric oxide in the central nervous system: Neuroprotection versus neurotoxicity. Nature Reviews Neuroscience, 8, 766–775. 10.1038/nrn2214

Calixto, A. V., Vandresen, N., De Nucci, G., Moreno, H., & Faria, M. S. (2001). Nitric oxide may underlie learned fear in the elevated T-maze. Brain Research Bulletin, 55(1), 37–42. 10.1016/S0361-9230(01)00480-4

Calixto, A. V, Duarte, F. S., Moraes, C. K. L., Faria, M. S., & Lima, T. C. M. De. (2008). Nitric oxide involvement and neural substrates of the conditioned and innate fear as evaluated in the T-maze test in rats. Behavioural Brain Research, 189, 341–349. 10.1016/j.bbr.2008.01.018

Çalışkan, G., & Stork, O. (2018). Hippocampal network oscillations as mediators of behavioural metaplasticity: Insights from emotional learning. Neurobiology of Learning and Memory, 154, 37–53. 10.1016/j.nlm.2018.02.022

Chen, Y.-J., Raman, G., Bodendiek, S., O’Donnell, M. E., & Wulff, H. (2011). The KCa3.1 Blocker TRAM-34 Reduces Infarction and Neurological Deficit in a Rat Model of Ischemia/Reperfusion Stroke. Journal of Cerebral Blood Flow & Metabolism, 31, 2363– 2374. 10.1038/jcbfm.2011.101

Cinelli, M. A., Do, H. T., Miley, G. P., & Silverman, R. B. (2020). Inducible nitric oxide synthase: Regulation, structure, and inhibition. Medicinal Research Reviews, 40(1), 158–189. 10.1002/med.21599

Cui, H., Hayashi, A., Sun, H.-S., Belmares, M. P., Cobey, C., Phan, T., Schweizer, J., Salter, M. W., Wang, Y. T., Tasker, R. A., Garman, D., Rabinowitz, J., Lu, P. S., & Tymianski, M. (2007). PDZ protein interactions underlying NMDA receptor-mediated excitotoxicity and neuroprotection by PSD-95 inhibitors. Journal of Neuroscience, 27, 9901–9915. 10.1523/JNEUROSCI.1464-07.2007

da Silva Chaves, S. N., Dutra Costa, B. P., Vidal Gomes, G. C., Lima-Maximino, M., Pacheco Rico, E., & Maximino, C. (2020). NOS-2 participates in the behavioral effects of ethanol withdrawal in zebrafish. Neuroscience Letters, 728, 134952. 10.1016/j.neulet.2020.134952

Daskalakis, N. P., Yehuda, R., & Diamond, D. M. (2013a). Animal models in translational studies of PTSD. Psychoneuroendocrinology, 38, 1895–1911. 10.1016/j.psyneuen.2013.06.006

Daskalakis, N. P., Yehuda, R., & Diamond, D. M. (2013b). Animal models in translational studies of PTSD. Psychoneuroendocrinology, 38, 1895–1911. 10.1016/j.psyneuen.2013.06.006

Dunn, R. W., Reed, T. A. W., Copeland, P. D., & Frye, C. A. (1998). The nitric oxide synthase inhibitor 7-nitroindazole displays enhanced anxiolytic efficacy without tolerance in rats following subchronic administration. Neuropharmacology, 37, 899–904. 10.1016/S0028-3908(98)00076-8

Edwards, T. M., & Rickard, N. S. (2007). New perspectives on the mechanisms through which nitric oxide may affect learning and memory processes. Neuroscience & Biobehavioral Reviews, 31, 413–425. 10.1016/j.neubiorev.2006.11.001

Garthwaite, J. (2008). Concepts of neural nitric oxide-mediated transmission. European Journal of Neuroscience, 27, 2783–2802. 10.1111/j.1460-9568.2008.06285.x

Gerlach, G. F., Schrader, L. N., & Wingert, R. A. (2011). Dissection of the Adult Zebrafish Kidney. Journal of Visualized Experiments: JoVE, 54, 2839. 10.3791/2839

Griess, J. P. (1864). On a new series of bodies in which nitrogen is substituted for hydrogen. Philosophical Transactions of the Royal Society, 154, 667–731.

Harvey, B. H. (2006). Adaptive plasticity during stress and depression and the role of glutamate-nitric oxide pathways. South African Psychiatry Review, 9, 132–139. 10.4314/ajpsy.v9i3.30214

Harvey, B. H., Bothma, T., Nel, A., Wegener, G., & Stein, D. J. (2005). Involvement of the NMDA receptor, NO-cyclic GMP and nuclear factor K-β in an animal model of repeated trauma. Human Psychopharmacology, 20, 367–373. 10.1002/hup.695

Harvey, B. H., Oosthuizen, F., Brand, L., Wegener, G., & Stein, D. J. (2004). Stress-restress evokes sustained iNOS activity and altered GABA levels and NMDA receptors in rat hippocampus. Psychopharmacology, 175, 494–502. 10.1007/s00213-004-1836-4

Herculano, A. M., Puty, B., Miranda, V., Lima, M. G., & Maximino, C. (2015). Interactions between serotonin and glutamate-nitric oxide pathways in zebrafish scototaxis. *Pharmacology*, Biochemistry & Behavior, 129, 97–104. 10.1016/j.pbb.2014.12.005

Jesuthasan, S. J., & Mathuru, A. S. (2008). The alarm response in zebrafish: Innate fear in a vertebrate genetic model. Journal of Neurogenetics, 22, 211–229. 10.1080/01677060802298475

Kaushal, V., Koeberle, P. D., Wang, Y., & Schlichter, L. C. (2007). The Ca2+-Activated K+ Channel KCNN4/KCa3.1 Contributes to Microglia Activation and Nitric Oxide-Dependent Neurodegeneration. Journal of Neuroscience, 27, 234–244. 10.1523/JNEUROSCI.3593-06.2007

Kinkel, M. D., Eames, S. C., Philipson, L. H., & Prince, V. E. (2010). Intraperitoneal injection into adult zebrafish. Journal of Visualized Experiments, 42, 2126. 10.3791/2126

Kornau, H.-C., Schenker, L. T., Kennedy, M. B., & Seeburg, P. H. (1995). Domain interaction between NMDA receptor subunits and the postsynaptic density protein PSD-95. Science, 269, 1737–1740. 10.1126/science.7569905

Lawford, B. R., Morris, C. P., Swagell, C. D., Hughes, I. P., Young, R. M., & Voisey, J. (2013). NOS1AP is associated with increased severity of PTSD and depression in untreated combat veterans. Journal of Affective Disorders, 147, 87–93. 10.1016/j.jad.2012.10.013

Lawrence, C. (2007). The husbandry of zebrafish (Danio rerio): A review. Aquaculture, 269(1), 1–20. 10.1016/j.aquaculture.2007.04.077

Lima, M. G., Silva, R. X. do C., Silva, S. de N. dos S., Rodrigues, L. do S. dos S., Oliveira, K. R. H. M., Batista, E. de J. O., Maximino, C., & Herculano, A. M. (2016). Time-dependent sensitization of stress responses in zebrafish: A putative model for post-traumatic stress disorder. Behavioural Processes, 128, 70–82. 10.1016/j.beproc.2016.04.009

Lima, M. G., Silva, S. de N. dos S., Silva, R. X. do C., Oliveira, K. R. H. M., Batista, E. de J. O., Maximino, C., & Herculano, A. M. (2015). Putative involvement of the nitrergic system on the consolidation, but not initiation, of behavioral sensitization after conspecific alarm substance in zebrafish. Pharmacology, Biochemistry & Behavior, 139, 127–133. 10.1016/j.pbb.2015.08.005

Lima-Maximino, M., Pyterson, M. P., do Carmo Silva, R. X., Gomes, G. C. V., Rocha, S. P., Herculano, A. M., Rosemberg, D. B., & Maximino, C. (2020). Phasic and tonic serotonin modulate alarm reactions and post-exposure behavior in zebrafish. Journal of Neurochemistry, 153, 495–509. 10.1111/jnc.14978

Luo, C., & Zhu, D. (2011). Research progress on neurobiology of neuronal nitric oxide synthase. Neuroscience Bulletin, 27, 23–35. 10.1007/s12264-011-1038-0

Łuszczki, J. J., Prystupa, A., Andres-Mach, M., Marzęda, E., & Florek-Łuszczki, M. (2013). Ivabradine (a hyperpolarization activated cyclic nucleotide-gated channel blocker) elevates the threshold for maximal electroshock-induced tonic seizures in mice. Pharmacological Reports, 65, 1407–1414. 10.1016/S1734-1140(13)71500-7

MacKenzie, G. M., Rose, S., Bland-Ward, P. A., Moore, P. K., Jenner, P., & Marsden, C. D. (1994). Time course of inhibition of brain nitric oxide synthase by 7-nitro indazole. NeuroReport, 5, 1993.

Maximino, C., do Carmo Silva, R. X., dos Santos Campos, K., de Oliveira, J. S., Rocha, S. P., Pyterson, M. P., dos Santos Souza, D. P., Feitosa, L. M., Ikeda, S. R., Pimentel, A. F. N. F. N., Ramos, P. N. F. N. F., Costa, B. P. D. P. D., Herculano, A. M., Rosemberg, D. B., Siqueira-Silva, D. H., & Lima-Maximino, M. G. (2019). Sensory ecology of ostariophysan alarm substances. Journal of Fish Biology, 95, 274–286. 10.1111/jfb.13844

Maximino, C., Marques de Brito, T., Dias, C. A. G. de M., Gouveia, A., & Morato, S. (2010). Scototaxis as anxiety-like behavior in fish. Nature Protocols, 5(2), Artigo 2. 10.1038/nprot.2009.225

Maximino, C., Meinerz, D. L., Fontana, B. D., Mezzomo, N. J., Stefanello, F. V., de S. Prestes, A., Batista, C. B., Rubin, M. A., Barbosa, N. V., Rocha, J. B. T., Lima, M. G., & Rosemberg, D. B. (2018). Extending the analysis of zebrafish behavioral endophenotypes for modeling psychiatric disorders: Fear conditioning to conspecific alarm response. Behavioural Processes, 149, 35–42. 10.1016/j.beproc.2018.01.020

Maximino, C., Puty, B., Benzecry, R., Araujo, J., Lima, M. G., Batista, E. de J. O., Oliveira, K. R. M., Crespo-López, M. E., Herculano, A. M., Araújo, J., Lima, M. G., de Jesus Oliveira Batista, E., Renata de Matos Oliveira, K., Crespo-Lopez, M. E., Herculano, A. M., Araujo, J., Lima, M. G., Batista, E. de J. O., Oliveira, K. R. M., … Herculano, A. M. (2013). Role of serotonin in zebrafish (*Danio rerio*) anxiety: Relationship with serotonin levels and effect of buspirone, WAY 100635, SB 224289, fluoxetine and para-chlorophenylalanine (pCPA) in two behavioral models. Neuropharmacology, 71, 83–97. 10.1016/j.neuropharm.2013.03.006

Mittal, A., & Kakkar, R. (2020). Nitric Oxide Synthases and Their Inhibitors: A Review. Letters in Drug Design & Discovery, 17, 228–252. 10.2174/1570180816666190222154457

Moraes, E. R. da S., Grisolia, A. B. A., Oliveira, K. R. M., Picanço-Diniz, D. L. W., Crespo-López, M. E., Maximino, C., Batista, E. de J. O., & Herculano, A. M. (2012). Determination of glutamate uptake by high performance liquid chromatography (HPLC) in preparations of retinal tissue. Journal of Chromatography B, 907, 1–6. 10.1016/j.jchromb.2012.07.027

Mutlu, O., Akar, F., Celikyurt, I. K., Tanyeri, P., Ulak, G., & Erden, F. (2015). 7-NI and ODQ Disturbs Memory in the Elevated plus Maze, Morris Water Maze, and Radial Arm Maze Tests in Mice. Drug Target Insights, 9, DTI.S23378. 10.4137/DTI.S23378

Parra, K. V., Adrian, J. C., & Gerlai, R. (2009). The synthetic substance hypoxanthine 3-N-oxide elicits alarm reactions in zebrafish (Danio rerio). Behavioural Brain Research, 205(2), 336–341. 10.1016/j.bbr.2009.06.037

Pokk, P., & Väli, M. (2002). The effects of the nitric oxide synthase inhibitors on the behaviour of small-platform-stressed mice in the plus-maze test. Progress in Neuro-Psychopharmacology & Biological Psychiatry, 26, 241–247.

Sadeghi, M. A., Hemmati, S., Nassireslami, E., Yousefi Zoshk, M., Hosseini, Y., Abbasian, K., & Chamanara, M. (2022). Targeting neuronal nitric oxide synthase and the nitrergic system in post-traumatic stress disorder. Psychopharmacology, 239, 3057–3082. 10.1007/s00213-022-06212-7

Schmidt, M. V., Abraham, W. C., Maroun, M., Stork, O., & Richter-Levin, G. (2013). Stress-induced metaplasticity: From synapses to behavior. Neuroscience, 250, 112–120. 10.1016/j.neuroscience.2013.06.059

Speedie, N., & Gerlai, R. (2008). Alarm substance induced behavioral responses in zebrafish (Danio rerio). Behavioural Brain Research, 188(1), 168–177. 10.1016/j.bbr.2007.10.031

Spiacci Jr, A., Kanamaru, F., Guimarães, F. S., & de Oliveira, R. M. W. (2008). Nitric oxide-mediated anxiolytic-like and antidepressant-like effects in animal models of anxiety and depression. Pharmacology, Biochemistry and Behavior, 88, 247–255. 10.1016/j.pbb.2007.08.008

Stam, R. (2007a). PTSD and stress sensitisation: A tale of brain and body. Part 1: Human studies. Neuroscience & Biobehavioral Reviews, 31, 530–557. 10.1016/j.neubiorev.2006.11.010

Stam, R. (2007b). PTSD and stress sensitisation: A tale of brain and body Part 2: Animal models. Neuroscience and Biobehavioral Reviews, 31(4), 558–584. 10.1016/j.neubiorev.2007.01.001

Stam, R., Bruijnzeel, A. W., & Wiegant, V. M. (2000). Long-lasting stress sensitisation. European Journal of Pharmacology, 405, 217–224. 10.1016/S0014-2999(00)00555-0

Theron, V., Harvey, B. H., Botha, T., Weinshenker, D., & Wolmarans, D. W. (2023). Life-threatening, high-intensity trauma- and context-dependent anxiety in zebrafish and its modulation by epinephrine. Hormones and Behavior, 153, 105376. 10.1016/j.yhbeh.2023.105376

Thomas, D. D., Heinecke, J. L., Ridnour, L. A., Cheng, R. Y., Kesarwala, A. H., Switzer, C. H., McVicar, D. W., Roberts, D. D., Glynn, S., Fukuto, J. M., Wink, D. A., & Miranda, K. M. (2015). Signaling and stress: The redox landscape in NOS2 biology. Free Radical Biology and Medicine, 87, 204–225. 10.1016/j.freeradbiomed.2015.06.002

Xu, X., Bazner, J., Qi, M., Johnson, E., & Freidhoff, R. (2003). The role of telencephalic NMDA receptors in avoidance learning in goldfish (*Carassius auratus*). Behavioral Neuroscience, 117, 548–554. 10.1037/0735-7044.117.3.548

Xu, X., Bentley, J., Miller, T., Zmolek, K., Kovaleinen, T., Goodman, E., & Foster, T. (2009). The role of telencephalic NO and cGMP in avoidance conditioning in goldfish (Carassius auratus). Behavioral Neuroscience, 614–623. 10.1037/a0015243

Xu, X., Scott-Scheiern, T., Kempker, L., & Simons, K. (2007a). Active avoidance conditioning in zebrafish (*Danio rerio*). Neurobiology of Learning and Memory, 87, 72–77. 10.1016/j.nlm.2006.06.002

Xu, X., Scott-Scheiern, T., Kempker, L., & Simons, K. (2007b). Active avoidance conditioning in zebrafish (Danio rerio). Neurobiology of Learning and Memory, 87, 72–77. 10.1016/j.nlm.2006.06.002

Yang, L., Wang, J., Wang, D., Hu, G., Liu, Z., Yan, D., Serikuly, N., Alpyshov, E. T., Demin, K. A., Strekalova, T., de Abreu, M. S., Song, C., & Kalueff, A. V. (2020). Delayed behavioral and genomic responses to acute combined stress in zebrafish, potentially relevant to PTSD and other stress-related disorders: Focus on neuroglia, neuroinflammation, apoptosis and epigenetic modulation. Behavioural Brain Research, 389, 112644. 10.1016/j.bbr.2020.112644

Yehuda, R., & Antelman, S. M. (1993). Criteria for rationally evaluating animal models of posttraumatic stress disorder. Biological Psychiatry, 33, 479–486. 10.1016/0006-3223(93)90001-T

